# The single-molecule accessibility landscape of newly replicated mammalian chromatin

**DOI:** 10.1101/2023.10.09.561582

**Authors:** Megan S Ostrowski, Marty G Yang, Colin P McNally, Nour J Abdulhay, Simai Wang, Elphège P Nora, Hani Goodarzi, Vijay Ramani

## Abstract

The higher-order structure of newly replicated (*i.e.* ‘nascent’) chromatin fibers remains poorly-resolved, limiting our understanding of how epigenomes are maintained across cell divisions. To address this, we present <u>R</u>eplication-<u>A</u>ware <u>S</u>ingle-molecule <u>A</u>ccessibility <u>M</u>apping (RASAM), a long-read sequencing method that nondestructively measures genome-wide replication-status and protein-DNA interactions simultaneously on intact chromatin templates. We report that individual human and mouse nascent chromatin fibers are ‘hyperaccessible’ compared to steady-state chromatin. This hyperaccessibility occurs at two, coupled length-scales: first, individual nucleosome core particles on nascent DNA exist as a mixture of partially-unwrapped nucleosomes and other subnucleosomal species; second, newly-replicated chromatin fibers are significantly enriched for irregularly-spaced nucleosomes on individual DNA molecules. Focusing on specific *cis*-regulatory elements (*e.g.* transcription factor binding sites; active transcription start sites [TSSs]), we discover unique modes by which nascent chromatin hyperaccessibility is resolved at the single-molecule level: at CCCTC-binding factor (CTCF) binding sites, CTCF and nascent nucleosomes compete for motifs on nascent chromatin fibers, resulting in quantitatively-reduced CTCF occupancy and motif accessibility post-replication; at active TSSs, high levels of steady-state chromatin accessibility are preserved, implying that nucleosome free regions (NFRs) are rapidly re-established behind the fork. Our study introduces a new paradigm for studying higher-order chromatin fiber organization behind the replication fork. More broadly, we uncover a unique organization of newly replicated chromatin that must be reset by active processes, providing a substrate for epigenetic reprogramming.

## INTRODUCTION

Nucleosomes are the linchpin of epigenetic inheritance. Histone-DNA interactions that form the nucleosome are disrupted during DNA replication, only to be quickly reformed through a mixture of histone recycling and *de novo* histone deposition behind the replication fork^1^. This assembly process is deterministic: chromatin structure established in early gap 1 (G1) phase is memorized, such that replicated chromatin in synthesis (S) and gap 2 (G2) phases retain transcriptional regulatory activity^2^. Mapping replication-coupled chromatin assembly is thus integral to understanding how gene expression programs are established within cell lineages and maintained across cell divisions.

Our knowledge of newly replicated chromatin structure has been built upon a series of pioneering biochemical experiments that revealed increased sensitivity of nascent nucleosomes to enzymatic digestion^3,4^. These approaches, however, are destructive by nature and do not provide insight into locus-specific chromatin assembly behind the replication fork. Genomic approaches combining classical nucleotide-analog labeling of nascent DNA^5^ with high-throughput sequencing of chromatin^6–12^ have increased resolution, but suffer from technical limitations associated with Illumina sequencing. For instance, micrococcal nuclease (MNase)-based approaches destroy accessible DNA, and Tn5-based approaches require multiple successive insertion events into DNA for library preparation; these activities potentially underestimate DNA accessibility behind the fork. Moreover, all existing approaches require population-averaging of short-read sequencing data, which has led to diverging biological interpretations. For instance, measurements of Tn5 hypersensitivity^9^ and MNase accessibility^13^ behind the replication fork in dividing murine embryonic stem cells (mESCs) offer opposing conclusions regarding the accessibility of motifs for the essential TF CCCTC-binding factor (CTCF) on nascent chromatin. Finally, genomic approaches that involve short-read sequencing fail to capture the connectivity of protein-DNA interactions on individual molecules, precluding an understanding of how chromatin fibers are repopulated by nucleosomes upon DNA replication. Ultimately, methodological limitations have left many questions regarding chromatin replication unanswered, including: how are nucleosomes structured and successively positioned on nascent DNA molecules? How is newly replicated chromatin structured in repetitive, typically ‘unmappable’ mammalian genomic sequence? How do sequence-specific transcription factors (TFs) bound in G1 re-engage with nascent chromatin fibers following replication? Answering these questions requires a genome-scale nascent chromatin profiling method capable of providing single-molecule information.

Inspired by single-molecule combing assays^14,15^ and recently-developed methods using Oxford Nanopore Technologies (ONT) sequencing^16–19^, we present a replication-aware long-read chromatin mapping technology that globally and non-destructively measures protein-DNA interactions on nascent chromatin fibers. We first demonstrate that long-read Pacific Biosciences (PacBio) sequencing can be used to natively detect the nucleotide analog BrdU, then further show that this can be combined with genome-wide single-molecule long-read chromatin accessibility mapping (as developed by us and others)^20–25^. Applying this method in human K562 cells and mESCs, we discover abundant heterogeneity in the structures of individual nascent chromatin fibers. We find that newly replicated chromatin is composed of partially-unwrapped nucleosomes and subnucleosomal species (*e.g.* hexasomes), that are arranged in a mixture of ordered (*i.e.* evenly-spaced) and disordered patterns on individual DNA molecules. These patterns contribute to a nearly genome-wide state of single-molecule DNA hyperaccessibility behind the replication fork that is not present in steady-state chromatin. Though hyperaccessibility occurs across different epigenomic domains and classes of genomic repeat sequence, we find that certain gene regulatory loci do not exhibit this hyperaccessible state; instead, they show chromatin accessibility levels that are more akin to those observed in mature chromatin. At CTCF motifs and actively-transcribed transcription start sites (TSSs), we demonstrate distinct replication-associated pathways for chromatin maturation: with CTCF sites, factor binding itself is reduced shortly after replication (<1 hr), CTCF motifs are usually nucleosome occluded or found in linker DNA, and sequence-dependent binding is required to re-establish single-molecule chromatin accessibility post-replication at cognate motifs; at active TSSs, high-levels of steady-state single-molecule accessibility are re-established within 10 minutes post-replication, corroborating recent evidence that pioneering rounds of transcription reset steady-state chromatin environments^9^. Our novel approach offers critical insight into the single-molecular competition between TFs and nucleosomes for regulatory DNA, while raising new mechanistic questions regarding chromatin maturation during the cell cycle.

## RESULTS

### Replication aware single-molecule accessibility mapping

Although computational methods for detecting halogenated nucleotides on the ONT platform exist^16–18^, methods for detecting BrdU on PacBio sequencers – which provide specific advantages over ONT sequencers^26^ – do not. We thus sought to engineer a predictive model that could classify single-molecule BrdU incorporation (*i.e.* BrdU^(+)^) using the measured kinetics of the PacBio sequencing polymerase^27^. Using a weighted mixture of PCR-derived-, murine, and human-genomic DNA samples containing dTTP or BrdUTP as training data, we trained a convolutional neural network (CNN; model schematic in **Supplementary Figure 1**) to predict BrdU^(+)^ from PacBio nucleotide incorporation kinetics and one-hot encoded DNA sequence, at the resolution of 500 basepair (bp) ‘tiles’ of double-stranded DNA (dsDNA). The CNN model generated a continuous prediction score (**Supplementary Figure 2A**), which could be used to accurately classify held-out positive and negative control tiles (AUC = 0.934; **Supplementary Figure 2B**). This demonstrates that PacBio sequencers can natively detect BrdU-containing DNA molecules.

We leveraged our model to simultaneously quantify single-molecule BrdU incorporation in dividing human and mouse cells (**Figure 1A**). We performed a series of experiments in which human (K562) or mouse (E14 mESC) cells were cultured in BrdU-containing media, footprinted using saturating amounts of the highly-active nonspecific adenine methyltransferase EcoGII^28^, and processed for high-fidelity (HiFi) PacBio sequencing on the Sequel II platform. To calibrate BrdU^(+)^ classification on long sequenced molecules with multiple 500 bp tiles, we determined an empirical false discovery rate by comparing the predicted fraction of BrdU^(+)^ molecules in labeled cells versus unlabeled control cells. This allowed us to determine classification cutoffs minimizing false positive BrdU^(+)^ detection (0.32% in K562; 0.21% in E14 control experiments; **Supplementary Figure 3A**). False positive rates were stable across multiple sequencing parameters, including AT-content (**Supplementary Figure 3B**), number of circular consensus sequencing (CCS) passes (**Supplementary Figure 3C**), and read length (**Supplementary Figure 3D**), demonstrating technical robustness of our approach. We emphasize that our classification strategy prioritizes specificity over sensitivity; this *in silico* design choice is analogous to short-read approaches that rely on biotin-enrichment or immunoprecipitation to select for analog-labeled molecules. Future computational tool development will address sensitivity limitations, such as nucleotide and strand-resolution of BrdU incorporation, associated with the proof-of-concept presented here.

**Figure 1:**
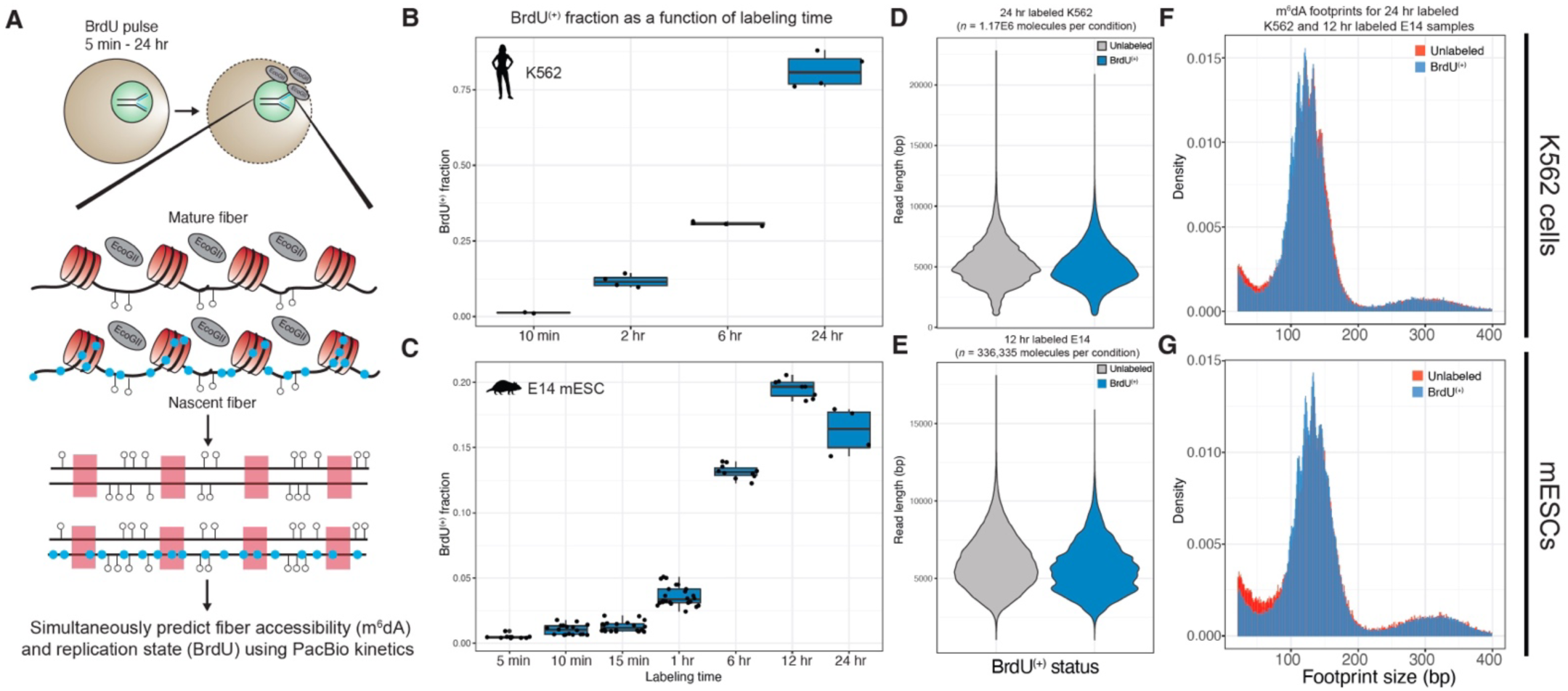
PacBio sequencing can simultaneously detect BrdUTP incorporation and m^6^dA-labeled chromatin accessibility on single DNA molecules from mammalian cells. **A.)** Schematic for the replication-<u>a</u>ware single-molecule <u>a</u>ccessibility <u>m</u>apping (RASAM) assay. Cultured cells are fed BrdU via culture media, which is incorporated into nascent DNA strands as BrdUTP. RASAM relies on the ability of PacBio to detect modified nucleotides through changes in PacBio sequencing polymerase kinetics to natively localize m^6^dA at near-base-pair resolution, and classify sequenced double-stranded DNA (dsDNA) on the basis of BrdUTP incorporation. **B.)** Boxplot representation of predicted BrdU^(+)^ fraction for K562 cells cultured in BrdU-containing media for 10 minutes to 24 hours. Individual points represent separate PacBio sequencing experiments. **C.)** As in **B.)** but for E14 murine embryonic stem cells (mESCs) cultured in BrdU for 5 minutes to 24 hours. **D.)** Violin plot for sequenced DNA fragment lengths for 24 hour-labeled K562 cells, separated by BrdU^(+)^ labeling status. **E.)** As in D.) but for 12 hour-labeled E14 mESCs. **F.)** Length distribution for predicted methyltransferase-inaccessible footprints from 24 hour-labeled K562 cells following the RASAM assay, colored by BrdU^(+)^ labeling status. Distributions largely overlap, indicating that BrdU^(+)^ labeling does not impact the ability to footprint nucleosomes using the RASAM assay. **G.)** As in **F.)** but for 12 hour-labeled E14 cells following the RASAM assay.

Having established cutoffs for specific BrdU^(+)^ classification, we examined the fraction of BrdU^(+)^ classified molecules as a function of labeling time in K562 and mESCs, from 5 minutes (min) to 24 hours (hr; **Figure 1B**). As expected, BrdU^(+)^ fraction reproducibly increased as a function of labeling time: K562 cells incorporated a minimum of 1.41% (1.15% in biological replicate) at 10 min, and a maximum of 81.4 ± 5.80% at 24 hr (mean ± standard deviation; *n =* 4 sequencing runs); mESCs incorporated a minimum of 0.488 ± 0.14% at 5 min, and a maximum of 19.5 ± 0.696% at 12 hr (*n =* 8 sequencing runs). In mESC data, we observed a slight dip in BrdU^(+)^ fraction in labeling experiments longer than one mES cell cycle, and speculate that the observed, reproducible differences in maximum BrdU^(+)^ fraction between K562 and mESC may be due to cell-type-specific differences in stable BrdU-incorporation rate. Technically, we also observed systematically-decreased BrdU^(+)^ fraction in longer molecules with fewer CCS passes, consistent with increased classification uncertainty for molecules with less kinetic data (**Supplementary Figure 4**).

We next assessed whether BrdU-incorporation impacted the ability to i.) sequence long, multi-kilobase DNA molecules, and ii.) accurately call methyltransferase-inaccessible footprints indicative of nucleosome positions. First, we visualized sequenced DNA fragment length distributions from BrdU^(+)^ molecules and unlabeled molecules from long-time point K562 (24 h; **Figure 1C**) and E14 (12 hr; **Figure 1D**) data. Observed sequenced fragment lengths differed only slightly, indicating that BrdU-incorporation does not substantially alter PacBio sequencer processivity. Second, we examined methylase-inaccessible ‘footprint’ distributions from long-labeling experiments for K562 (**Figure 1E**) and E14 (**Figure 1F**). These distributions were also highly consistent, demonstrating strong enrichment for mononucleosome-sized footprints on BrdU^(+)^ and unlabeled molecules alike. Together, these results indicate that PacBio sequencing can coincidentally measure BrdU-incorporation and chromatin accessibility on long individual DNA molecules purified from mammalian cells. Hereafter, we refer to this approach as <u>R</u>eplication <u>A</u>ware <u>S</u>ingle-molecule <u>A</u>ccessibility <u>M</u>apping, or RASAM.

### Nascent mammalian chromatin fibers are hyperaccessible

In total, we generated RASAM data in biological duplicate or triplicate across four different labeling time points (10 min, 2 hr, 6 hr, 24 hr) for K562 cells (28.1 Gb), and six different labeling time points (5 min, 10 min, 1 hr, 6 hr, 12 hr, and 24 hr) for E14 mESCs (78.1 Gb; library summary statistics in **Supplemental Table 1**). To leverage this unique resource, we focused our analyses on labeling time points between 10 min and 24 / 12 hr for K562s / mESCs, respectively; we selected these timepoints to balance sampling of multiple labeling durations with sequencing a reasonable number of BrdU^(+)^ molecules at short labeling time points.

We began by quantifying differences in single-molecule chromatin accessibility in BrdU^(+)^ versus unlabeled molecules across labeling times. We computed the average accessibility of sampled BrdU^(+)^ and unlabeled molecules, which we used to calculate an ‘accessibility fold change’: the fold-difference in mean accessibility between BrdU^(+)^ and unlabeled molecules for each sequenced sample (**Figure 2A**). At early labeling times (*i.e.* 10 min – 6 hr), when BrdU^(+)^ fibers most likely represent nascent chromatin, we were surprised to observe high accessibility fold changes (**Figure 2B,C**). These increases ranged in effect-size: in K562s, from 37.6%, 48.7%, and 45.4% at 10 min, 2 hr, and 6 hr, respectively; in mESC, from 23.2%, 20.1%, and 23.8% at 10 min, 15 min, and 1 hr, respectively. These effects were reproducible (see individual points in **Figure 2B,C**; replicate values tabulated in **Supplemental Table 2**). Notably, these effects were also labeling-time dependent (effect sizes of 3.32% and 3.49% in replicate 24 hr K562 experiments; 2.62% and −0.346% in replicate 12 hr E14 experiments), consistent with long labeling-time BrdU^(+)^ fibers more closely matching steady-state fibers. Together, these analyses suggest that individual nascent chromatin fibers in dividing human and mouse cells are more accessible than steady-state fibers.

**Figure 2:**
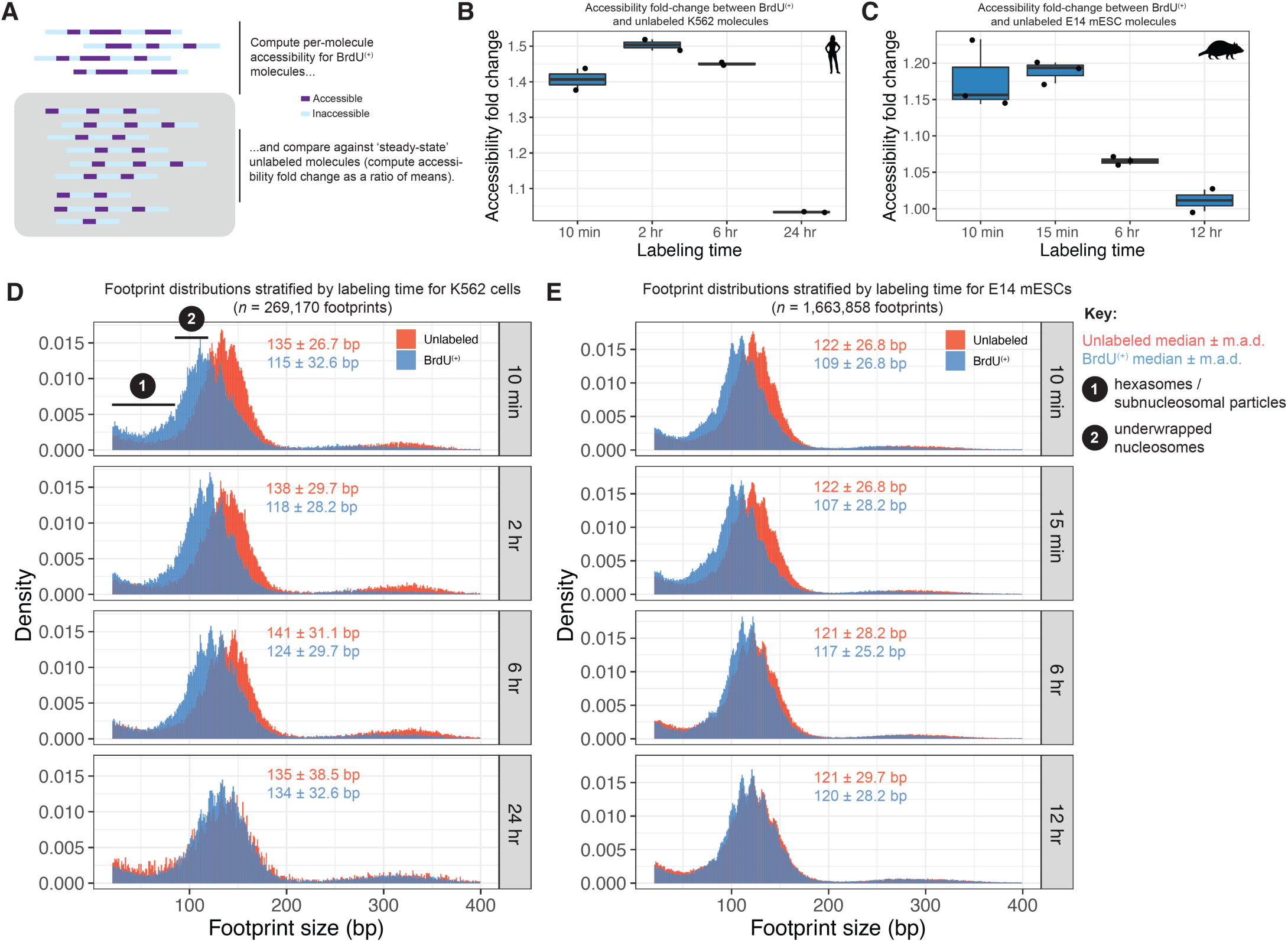
RASAM reveals single-molecule DNA hyperaccessibility on newly-replicated mammalian chromatin fibers. **A.)** Schematic of computational analysis to assess fold-differences between single-molecule DNA accessibility on BrdU^(+)^ versus unlabeled molecules. We compute the mean accessibility of each individual molecule, and then compare the mean BrdU^(+)^ single-molecule accessibility to the mean unlabeled single-molecule accessibility for different labeling times. **B.)** Box plot of the ratio of means (*i.e.* ‘accessibility fold change’) for K562 across four different labeling times. Box plot represents measurements from *n* = 2 biological replicates (*i.e.* separatae RASAM experiments). **C.)** Box plot of the accessibility fold change for E14 across four different labeling times. Box plots represent measurements from *n* = 3 biological replicates for 10 min., 15 min., and 6 hour labeling times, and *n* = 2 biological replicates for 12 hour labeling time. **D.)** Histogram of called methyltransferase-inaccessible footprints (*i.e.* ‘footprints’) for K562 cells across four labeling times. Footprints from BrdU^(+)^ fibers are colored in blue, while footprints for unlabeled fibers are colored in red. Median and median absolute deviation (m.a.d.) for each distribution shown in matching colors. Colored numbers highlight footprints with sizes of 1.) hexasomes and smaller, subnucleosomal particles, and 2.) ‘underwrapped’ nucleosome core particles. **E.)** Histogram of called footprints for E14 cells across four labeling times; coloring as in (**D**).

What could account for systematically-increased per-molecule accessibility in nascent chromatin? The primary constituent of chromatin is the octameric nucleosome, which protects ∼147 bp of DNA. As previously demonstrated by our group *in vitro* and *in vivo*, this footprint is shorter in nondestructive m^6^dA-footprinting experiments^20,25^, likely due to dynamic DNA breathing at nucleosome edges^25^. To investigate how nucleosomal protection of DNA differs on nascent chromatin, we plotted the distribution of observed footprint sizes for BrdU^(+)^ molecules at various time points, for K562 (**Figure 2D**) and mESCs (**Figure 2E**). In both cases, we observed clear, labeling-time dependent shifts in the amount of DNA protected by individual nucleosomes on BrdU^(+)^ versus steady-state molecules, indicative of DNA unwrapping from the nucleosome core particle. In 10 min K562 RASAM data, steady-state K562 nucleosomes protected 135 ± 26.7 bp (median ± median absolute deviation for footprints sized between 20 and 400 bp), while BrdU^(+)^ nascent footprints protected 115 ± 26.7 bp. In 10 min E14 RASAM data, steady-state nucleosomes protected less DNA than in K562 (122 ± 26.8 bp), and nascent footprints were even shorter (109 ± 26.8 bp). These labeling-time-specific distributions (**Figure 2D,E**; **inset**) imply that nascent nucleosomes are partially-unwrapped (possibly during an assembly process), and that differences in the extent of nucleosome wrapping in steady-state chromatin also contribute to observed cell-type-specific differences in accessibility fold change for nascent nucleosomes.

Footprint sizes can be used to infer the wrapping states of subnucleosomal particles such as hexasomes (*i.e.* nucleosomes that have lost a single H2A/H2B dimer)^29–31^. Visual inspection of BrdU^(+)^ footprint size distributions across labeling times revealed multiple features suggesting that nascent chromatin fibers are populated not only by unwrapped nucleosomes, but also subnucleosomal particles (see **Figure 2D, labels**). A relative enrichment of footprints ∼80 bp and below are consistent with enrichment of hexasomes and smaller subnucleosomal particles (*e.g.* tetrasomes devoid of both H2A/H2B dimers) in nascent DNA. We conclude that newly-replicated human and mouse chromatin fibers exist in a state of increased single-molecule accessibility – a phenomenon we term “nascent chromatin hyperaccessibility.” Furthermore, we conclude that nascent chromatin hyperaccessibility is due to systematic under-wrapping of constituent nucleosomal particles and the presence of subnucleosomal particles on newly replicated chromatin fibers.

### Nascent chromatin fibers are rapidly remodeled and enriched for disordered nucleosomal arrays

Nascent DNA hyperaccessibility may also arise from specific intramolecular patterns of nucleosomes / subnucleosomes arranged into nucleosomal arrays, a feature that cannot readily be surveyed by short-read sequencing measurements that destroy the chromatin fiber. To determine single-molecule nucleosome patterning on nascent chromatin, we analyzed RASAM data at the level of single-molecule nucleosome repeat length (NRL) and nucleosome regularity. We applied a single-molecule analytical strategy developed previously by our group (**Figure 3A**)^20,25^: briefly, we computed single-molecule autocorrelograms from BrdU^(+)^ molecules in K562 (labeling times ranging from 10 min to 24 hr) and mESC (labeling times ranging from 10 min to 12 hr); these autocorrelograms capture the regularity and average spacing of nucleosomes on individual chromatin fibers. We then employed unbiased community detection on *k*-nearest-neighbors (*k*NN) graphs (‘leiden’ clustering)^32^ of individual autocorrelograms, to group individual molecules into clusters we refer to as chromatin ‘fiber types.’ Finally, we carried out enrichment tests to ascertain differences in the relative proportion of specific fiber types (between nascent and steady-state molecules) as a function of labeling time.

**Figure 3:**
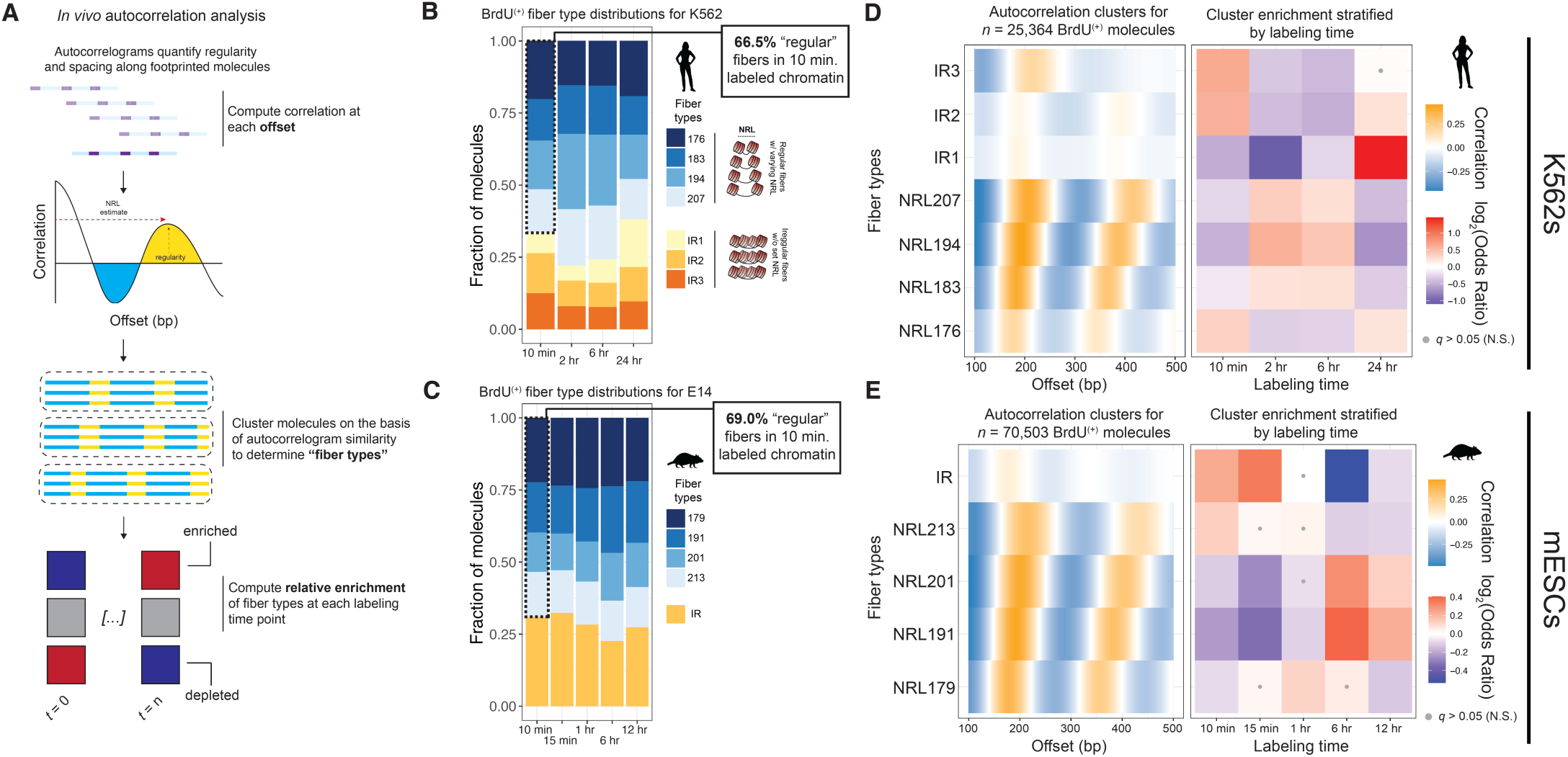
Global analysis of single-molecule nucleosome spacing in nascent and maturing chromatin. **A.)** Schematic of single-molecule autocorrelation analysis used for this study. As previously, we used single-molecule autocorrelograms to quantify the spacing and regularity of individual footprinted molecules in RASAM data. We first compute the autocorrelation function for the first thousand nucleotides of every sequenced molecule; we then use leiden community detection to cluster these single-molecule autocorrelograms to determine “fiber types” with shared patterns of nucleosome spacing and regularity; finally we perform enrichment tests to determine statistically-significant enrichment and depletion of specific fiber types in BrdU^(+)^ molecules across the full range of labeling times. **B.)** Stacked bar chart of fiber type representation for K562 cells. 66.5% of BrdU(+) fibers following 10 min labeling exhibit regularly spaced nucleosomes on single DNA molecules. **C.)** Stacked bar chart of fiber type representation for E14 cells. 69.0% of BrdU(+) fibers following 10 min. labeling exhibit regularly spaced nucleosomes on single DNA molecules. **D.)** Left: visualization of average autocorrelation of molecules falling into one of seven different fiber types for K562 cells; right: heatmap of log2(odds ratios) (*i.e.* ‘effect sizes’) capturing enrichment (red) or depletion (blue) of specific fiber types at each labeling time. Tests that are not significant with a false discovery rate of 5% (*q* > 0.05) are marked with a grey dot. **E.)** As in (**D**) but for E14 mESCs.

Applying this pipeline to RASAM data from K562 and mESCs, we obtained seven distinct fiber types for K562s and five fiber types for mESCs. These fiber types were made up of regularly-spaced clusters of varying NRL (176 bp – 207 bp in K562; 179 bp – 213 bp in mESC), and ‘irregular’ clusters where nucleosomes are unevenly spaced on individual templates (3 irregular clusters [IR1 – IR3] in K562; 1 irregular IR cluster in mESC). We then visualized distributions of these fiber types as stacked barplots stratified by labeling time, for K562 (**Figure 3B**) and mESC (**Figure 3C**). In both cell types, we observed a striking percentage of regular BrdU^(+)^ fibers (66.5% in K562; 69.0% in mESC), even at 10 min of labeling. These distributions imply that chromatin remodeling rapidly sets evenly spaced nucleosomes and subnucleosomes within 10 minutes post-replication.

We next tested the hypothesis that specific fiber types are enriched or depleted in nascent chromatin. For each fiber type (**Figure 3D,E**; **left**) and labeling time, we performed a series of Fisher’s exact enrichment tests to assess statistically-significant enrichment, which we visualized as a heatmap of effect sizes (**Figure 3D,E**; **right**). In K562 cells, we observed statistically-significant enrichment of one regular fiber type with a short repeat length (NRL176) and two irregular fiber types (IR2, IR3) at 10 min labeling (NRL176 Odds Ratio [OR] = 1.27, Storey’s *q*-value [*q*] = 3.33E-10; IR2 OR = 1.50, *q* = 2.70E-19; IR3 OR = 1.55, *q* = 5.55E-20). In mESCs, we observed significant enrichment of one regular fiber type with a long NRL of 213, as well as the single irregular fiber type (IR) at 10 min labeling (NRL213 OR = 1.10, *q* = 1.06E-3; IR OR = 1.17, *q* = 2.56E-14), as well as significant enrichment of irregular fibers at 15 min labeling (IR OR = 1.27, *q* = 1.17E-31). Collectively, these analyses demonstrate that nascent mammalian chromatin fibers are enriched for irregular, disordered nucleosome arrays.

Nucleosome positioning *in vitro* is strongly biased by primary DNA sequence^33^, though *in vivo* positioning is significantly less sequence-constrained due to the effects of ATP-dependent chromatin remodelers^34^. We hypothesized that primary sequence may provide more of a constraint on nucleosome positions on irregular, nascent nucleosome arrays than steady-state chromatin, possibly due to RASAM sampling individual fibers prior to replication-associated chromatin remodeling and maturation. To test this hypothesis, we computed Pearson’s product moment correlations (Pearson’s *r*) between accessibility and windowed AT-dinucleotide content for each individual molecule. This was done with the expectation that increased single-molecule correlation denotes increased sequence-constraint on observed single-molecule nucleosome positions. We performed this analysis specifically on fiber types enriched at early labeling times (irregular fibers in K562; irregular and long NRL fibers in mESC), and visualized results as a boxplot of single-molecule Pearson’s *r* values (**Supplementary Figure 5**). For K562, we observed significantly higher correlations (between accessibility and AT-content) for nascent irregular fibers compared to steady-state fibers (student’s *t*-test *p* = 1.54E-12); by comparison, we observed no significant difference in nascent regular fibers compared to steady-state regular fibers (*p* = 0.21). We observed similarly significant (though smaller effect size) trends in mESC: for both irregular and long NRL fibers, as well as short NRL fibers, we observed significantly higher Pearson’s *r* values in nascent versus steady-state fibers (*p* =1.12E-14 and *p* =3.65E-10, respectively). Taken as a whole, these analyses demonstrate that: i.) chromatin remodeling evenly spaces unwrapped nucleosomes and subnucleosomes within 10 min following replication; ii.) that nucleosomal arrays assembled on nascent DNA are statistically enriched for irregularly-spaced nucleosomes; and, finally iii.) that nucleosome positions on irregular nascent fibers are more constrained by primary DNA sequence than those on mature fibers.

### Epigenome-wide nascent DNA hyperaccessibility

The epigenome is functionally compartmentalized into active and silent domains. We wondered whether under-wrapped nucleosomes were a feature specific to fibers from particular epigenomic domains. To test this, we focused analyses on a mixture of “mappable” epigenomic domains (H3K4me3 [active promoters], H3K4me1 [active enhancers], H3K36me3 [actively transcribed gene bodies], H3K27me3 [facultative repression], H3K9me3 [constitutive repression]), and unmappable regions uniquely accessible by long-read technology (murine major satellite [MajSat], murine minor satellite [MinSat], murine telomeric sequence, and murine ribosomal DNA [rDNA]). We performed these and all subsequent analyses on our mESC RASAM dataset, using mouse ENCODE^35^ and other published epigenomic datasets^36–38^ as references.

We computationally filtered for BrdU^(+)^ and unlabeled reads via reference genome mapping (for mappable regions and mouse rDNA), or by directly scanning PacBio CCS reads for matches to known repetitive sequence motifs (for MajSat, MinSat, and telomeric sequence), then visualized histograms of footprint sizes for filtered reads stratified by labeling time (**Figure 4**; statistics tabulated in **Supplementary Table 3**). Steady-state footprint size distributions were largely consistent across domains (ranging from 120 ± 19.0 bp in H3K4me1 domains to 130 ± 19.0 bp in MinSat sequence), and nascent (10 min – 1 h) footprint size distributions all displayed strongly significantly-lower median values across analyzed domains (*e.g.* H3K27me3 Mann-Whitney one-sided *p* = 2.39E-96; all one-sided Mann-Whitney *p* values tabulated in **Supplementary Table 3**). We conclude from these analyses that nascent chromatin hyperaccessibility is a general feature of the genome, including loci present in satellite, telomeric, and rDNA sequences not readily surveyable by Illumina sequencing. Given that this state is present in all epigenomic domains surveyed (each with a unique complement of *trans-*acting factors), this hyperaccessible state is likely a general property of loci that have been recently replicated.

**Figure 4:**
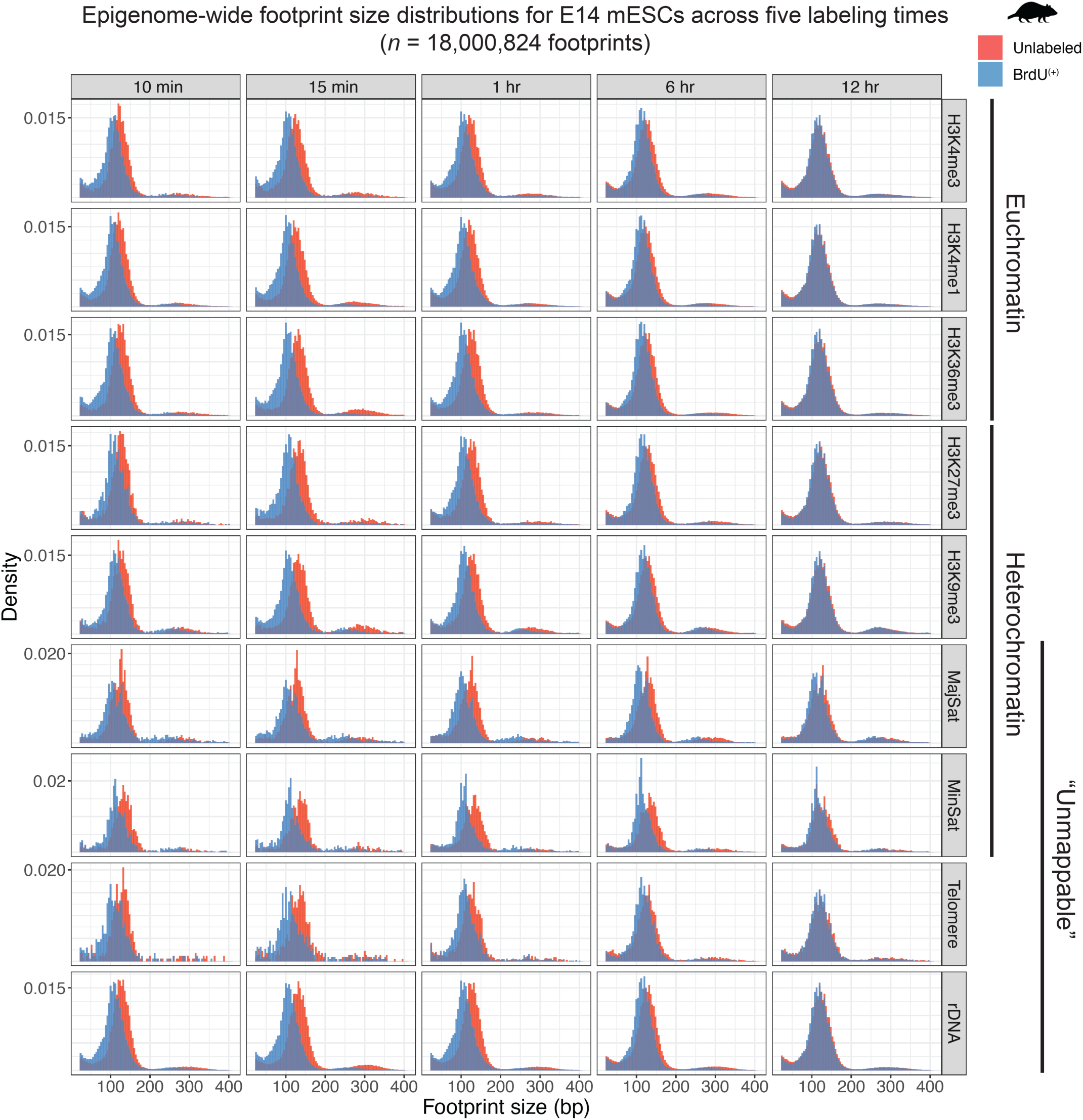
Patterns of single-molecule DNA hyperaccessibility are shared across epigenomic domains in E14 mESCs. Histograms of footprint sizes across five different labeling time points in E14 mESCs, for mappable euchromatic domains (H3K4me3, H3K4me1, H3K36me3), mappable heterochromatic domains (H3K27me3, H3K9me3), unmappable heterochromatic domains (murine major satellite, murine minor satellite, murine telomeric sequence), and typically unmappable ribosomal DNA (rDNA). BrdU^(+)^ nucleosomes are colored in blue, and unlabeled nucleosomes are colored in red.

### Nascent chromatin fiber structure at transcription factor binding motifs and gene promoters

Nascent chromatin hyperaccessibility raises an important question: how is regulatory specificity at cell-type specific *cis*-regulatory sequences (*e.g.* TF binding sites, TSSs) achieved given pervasive nucleosome unwrapping and globally increased DNA accessibility? We broadly envisioned four possible patterns by which individual chromatin fibers could be reformed at *cis*-regulatory regions (visualized in **Supplementary Figure 6**): first, chromatin fibers composed of mature nucleosomes could be repopulated with partially-unwrapped nucleosomes (nascent chromatin hyperaccessibility; **Supplementary Figure 6A**); second, chromatin fibers composed of mature nucleosomes may not be rapidly repopulated with nucleosomes, allowing TFs to bind their recognition motifs (another form of nascent chromatin hyperaccessibility; **Supplementary Figure 6B**); third, chromatin fibers devoid of nucleosomes could be repopulated with partially-unwrapped nucleosomes (nascent chromatin hypoaccessibility; **Supplementary Figure 6C**); or fourth, chromatin fibers devoid of nucleosomes could remain nucleosome free (nascent chromatin accessibility remains the same; **Supplementary Figure 6D**). To distinguish between and quantify how often these single-molecule states respectively occur at *cis-*regulatory motifs, we examined single-molecule nascent chromatin accessibility at ChIP-seq-backed binding sites (for CTCF, NRF1, REST, and SOX2) and active transcriptional start sites (TSSs).

For each factor, we calculated accessibility fold change for a 150 bp window surrounding motif occurrences within BrdU^(+)^ fibers and a matched number of randomly sampled unlabeled fibers, to quantify differences between nascent and steady-state accessibility. Given the sparsity of data at short labeling times, we controlled for random variation in this estimate by bootstrapping a distribution of accessibility fold change measurements per biological replicate per factor, such that each value represented a fold-change calculation between mean BrdU^(+)^ signal and signal from an equivalent number of sampled unlabeled molecules (**Supplementary Figure 7)**. We observed significantly lower accessibility fold changes (10 min *p* = 0.00133; 15 min *p* < 0.001; 60 min *p* < 0.001; one-sided permutation tests) for *bona fide* CTCF binding sites, compared to random negative control motifs (**Supplementary Figure 7A**); this trend was consistent (though varying in significance) across other tested factors and times (**Supplementary Figure 7B**), but not at TSSs (comparison of top 80% expressed TSS in mESC vs. bottom 20%; 10 min *p* = 0.117; 15 min *p* = 0.157; 60 min *p* = 0.333; **Supplementary Figure 7C**).

To further investigate this relationship at TF-binding sites, we visualized RASAM data centered at CTCF binding sites (**Figure 5A**), as line plots of average single-molecule accessibility for BrdU^(+)^ (blue) and unlabeled (red) fibers separated by labeling time. To account for data sparsity (all fiber counts in **Figure 5A,B**; **inset**), we plotted the average signal for unlabeled fibers as the mean ± standard deviation (gray ribbon) for *n* = 100 sampled unlabeled fiber equivalents. Examining unlabeled signal on these plots, we observed patterns broadly consistent with expected chromatin organization at CTCF sites and highly expressed TSSs. At CTCF sites, we observed well-phased nucleosomes characteristic of CTCF motifs (**Figure 5A**; **feature 1**)^39^, as well as a dip in signal representative of molecules where CTCF was bound during footprinting (**Figure 5A**; **feature 2**). Emergence of this feature as early as 10 min following replication is consistent with published nascent MNase footprinting data from mESCs^13^, while the observation of matched or decreased accessibility surrounding the CTCF motif at early time points suggests that nascent fibers are more often nucleosome-occluded compared with those at steady-state, consistent with repli-ATAC-seq measurements^9^. Identical analyses performed at NRF1 (**Supplementary Figure 9A**), REST (**Supplementary Figure 9C**), and SOX2 (**Supplementary Figure 9E**) demonstrate distinct, factor-specific patterns not seen in matched negative control motifs (**Supplementary Figure 9B,D,F**), suggesting that this reduction in hyperaccessibility is conserved across multiple TF families. Importantly, these patterns were highly-specific to bound CTCF sites; matched visualization of average single-molecule accessibility at control sites (**Supplementary Figure 8A**) demonstrated elevated signal in BrdU^(+)^ fibers consistent with nascent DNA hyperaccessibility. Altogether, these data indicate that the nascent chromatin hyperaccessibility observed genome-wide is restrained at TF-bound motifs, and not at randomly sampled motifs for these same factors.

**Figure 5:**
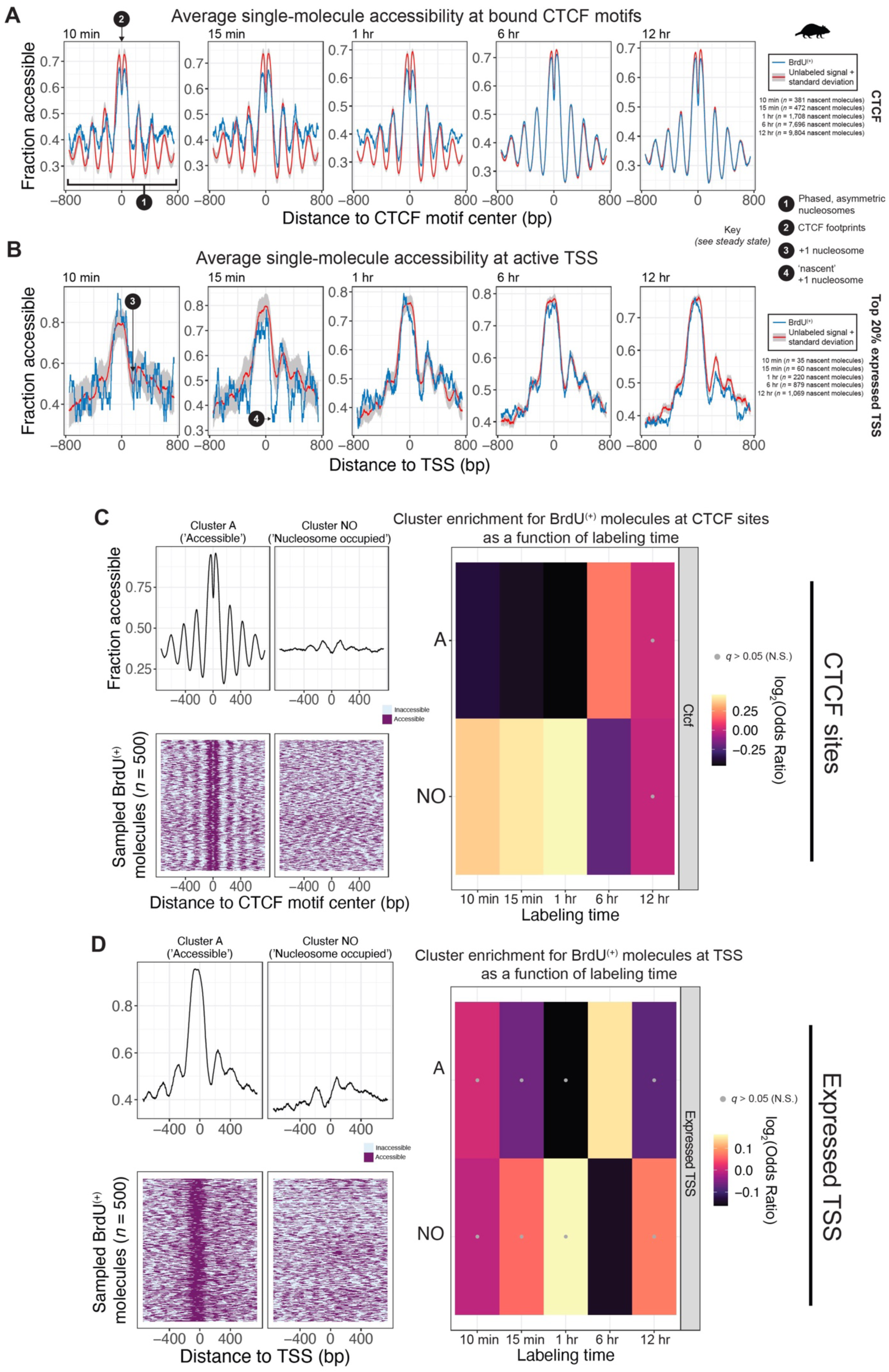
Patterns of CTCF motif and TSS accessibility on nascent chromatin fibers. **A.)** Line plot visualization of average single-molecule accessibility of BrdU^(+)^ (blue) and unlabeled (red) fibers centered at ChIP-seq backed CTCF motifs, stratified by labeling time. Grey ribbons around unlabeled signal represent the standard deviation of mean accessibility calculated from *n* = 100 random samples of an equivalent number of unlabeled molecules, compared to BrdU^(+)^ molecules. **Inset:** tabulated BrdU^(+)^ fiber counts for each labeling time point; key describes 1.) asymmetrically phased nucleosomes surrounding the CTCF motif (in steady state signal), as expected and 2.) position of +1 nucleosome in steady-state signal, downstream of highly expressed TSSs. **B.)** As in (**A**), but for fibers overlapping with a TSS for the top 20% highest expressed genes in mESC. **C.) Top:** line plot representations of two clusters of nascent, bound CTCF motif-containing molecules, which broadly represent a motif accessible cluster (*i.e.* ‘A’), and a nucleosome-occupied cluster (*i.e.* ‘NO’). Dip in signal at CTCF motif represents cases where CTCF itself was footprinted on chromatin. **Bottom:** Sampled single-molecules underling the averages shown as lineplots for each cluster. Each line represents an 1500 nucleotides extracting from an individual molecule, centered at the CTCF motif. **Right:** Heatmap representation of enrichment (yellow) or depletion (black) of clusters A and NO across labeling times. Grey dots mark Fisher’s exact tests where *q* > 0.05 (not significant). **D.)** As in (**C**) but for expressed TSSs (note unlike in (**B**), now including the top 80% of expressed genes in mESC). Fisher’s exact tests all demonstrate effect sizes close to 1, indicating no significant difference in cluster distribution between BrdU^(+)^ and unlabeled molecules across most labeling times.

In contrast, at highly expressed TSSs we observed high accessibility upstream of the TSS, suggestive of nucleosome free regions (NFRs) on these molecules. We also observed patterns indicative of a well-positioned +1 nucleosome downstream of the TSS (**Figure 5B**, **feature 3**) that is restored in a directional manner as early as 15 min post-replication (**Figure 5B**; **feature 4**). This rapid restoration of chromatin accessibility patterns at TSS runs counter to previously published results; we propose that this discrepancy is owed to differences between nondestructively measuring chromatin accessibility through methyltransferases, compared to either destructive measurement of chromatin through MNase, or Tn5 insertion and sequencing. Rapid restoration of the NFR and positioning of the +1 nucleosome was less prevalent at control loci (in this case, TSS for lowly-expressed genes in mESCs; **Supplementary Figure 8B**), consistent with a role for transcription in re-establishing TSS architecture at critical cell-type-specific genes.

To determine how single-molecule chromatin accessibility is regulated at these loci and distinguish between the models illustrated in **Supplementary Figure 6**, we computationally classified accessibility patterns of CTCF binding sites and TSSs at single-molecule resolution. We aggregated all BrdU^(+)^ fibers underlying blue plotted averages, and subjected these molecules to leiden clustering, resulting in clusters for CTCF and TSS-containing molecules. We visualized the signal of these clusters in two ways: through line plots of the average signal for molecules within each cluster (**Figure 5C,D**; **top left**), and as individual, sampled molecules colored by single-molecule accessibility (**Figure 5C,D**; **bottom left**). For both CTCF sites and active TSSs, leiden clustering detected two distinct classes of accessibility; we termed these ‘accessible’ (A), for molecules that demonstrated focal accessibility of either the CTCF site or TSS, and ‘nucleosome occupied’ (NO), for molecules that demonstrated reduced accessibility owing to the stochastic positioning of a nucleosome over the CTCF site or TSS. Having classified molecules into one of these two states, we performed enrichment tests to assess significant enrichment or depletion of these states across labeling times (**Figure 5C,D right**). For CTCF, we noticed a striking and significant depletion of the A state in short labeling time molecules (10 min OR = 0.776, *q* = 3.92E-2; 15 min OR = 0.750, *q* = 7.25E-3; 1 hr OR = 0.728; *q* = 8.94E-9); this suggests that though CTCF does bind nascent chromatin, CTCF occupancy does not return to steady-state levels until after 1 hr. For actively transcribed TSSs, conversely, we observed a different pattern: states A and NO were evenly represented across all labeling times, as evidenced by low, insignificant effect sizes (State A 10 min OR = 1.005, *q =* 1.00; 15 min OR = 0.957, *q* = 0.983; 1 hr OR = 0.893, *q* = 0.193); these data suggest that TSS accessibility is programmed very rapidly – at least within 10 min following replication. Collectively, this suggests that different regulatory sequences harbor distinct single-fiber chromatin maturation dynamics: first, CTCF molecules compete with nascent nucleosomes for the same motifs, resulting in a quantitative reduction of CTCF occupancy and motif accessibility behind the replication fork (consistent with the pattern illustrated in **Supplementary Figure 6C**); second, TSSs retain their single-molecule state distributions immediately following replication, suggesting that the NFRs generated by the basal transcriptional machinery and associated regulatory factors are either retained or very rapidly re-established post-replication (consistent with the pattern illustrated in **Supplementary Figure 6D**).

### CTCF directly instructs nascent chromatin fiber structure

Our results for CTCF and other TFs imply that factor binding might directly regulate nascent chromatin hyperaccessibility. To test this model, we combined RASAM profiling with chemical degradation of CTCF in genetically-engineered E14 mESCs harboring the auxin degron system^40,41^. Our experimental schematic is illustrated in **Figure 6A**: following 6 hours of auxin treatment (Western blot validation of near-complete CTCF degradation shown in **Supplementary Figure 10**), we performed three different RASAM experiments in biological duplicate, labeling cells for 10 min, 15 min, or 1 hr, to address whether CTCF is acutely required shortly before and after replication fork passage for re-establishing accessibility patterns at their motifs.

**Figure 6:**
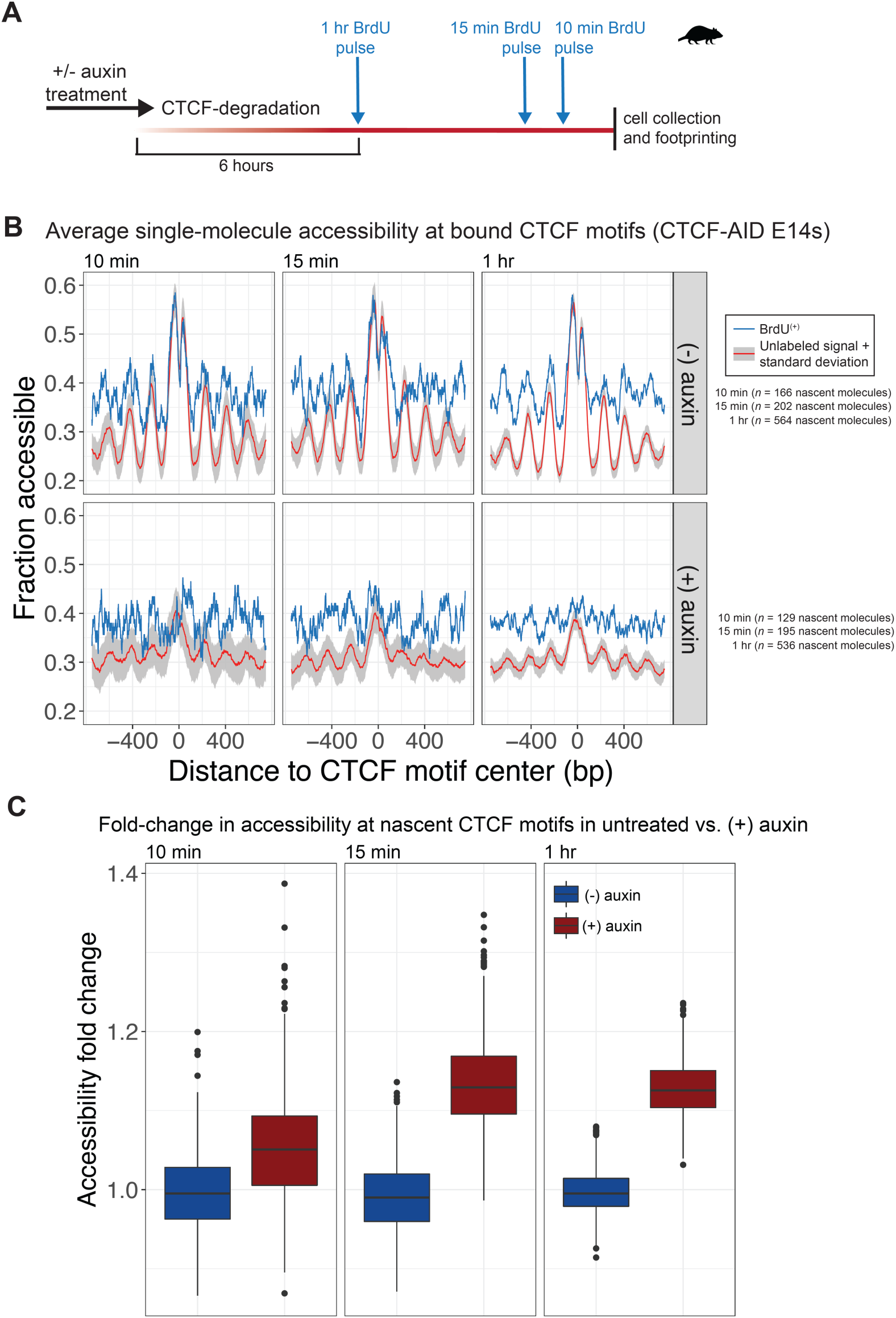
Acute CTCF degradation reveals that CTCF directly impacts DNA hyperaccessibility at bound motifs. **A.)** Schematic of experiment combining RASAM profiling with chemical degradation of the TF CTCF. **B.)** Average of single-molecule accessibility centered at ChIP-backed CTCF motifs for CTCF-AID E14 cells untreated (**top**) and treated with auxin (**bottom**). **C.)** Box plots of fold change accessibility estimates between BrdU^(+)^ and unlabeled molecules at CTCF motifs for untreated (blue) and treated (red) CTCF-AID E14 cells, stratified by labeling time. Box plot distributions represent the median and interquartile range for accessibility fold changes calculated from the average accessibility of BrdU^(+)^ molecules divided by *n* = 1,000 randomly sampled, equivalent numbers of unlabeled molecules per biological replicate (*n* = 2,000 values per boxplot).

We first performed line plot analyses at CTCF motifs for each experimental condition (**Figure 6B**). Average single-molecule accessibility patterns for BrdU^(+)^ and unlabeled molecules from (-) auxin cells resembled those of wildtype mESCs, consistent with previous reports that degron-tagging does not substantially alter CTCF occupancy^41^. Following CTCF degradation, however, we observed a striking increase in single-molecule accessibility on nascent chromatin compared to steady-state fibers. We quantified this difference by repeating the accessibility fold change analysis between BrdU^(+)^ and unlabeled fibers (as above; **Figure 6C**). CTCF degradation restored nascent chromatin hyperaccessibility at these motifs; accessibility fold changes were significantly higher in (+) auxin versus (-) auxin samples at 15 min and 60 min (15 min *p* = 0.001; 60 min *p* < 0.001, one-sided permutation test). This fold change in accessibility is roughly equivalent to the genome-wide difference observed between nascent and steady-state fibers at comparable early timepoints, suggesting that CTCF binding directly regulates nascent chromatin hyperaccessibility. Our results are consistent with the following model: upon replication, loci that harbor CTCF motifs are repopulated through a competition between nascent nucleosomes and CTCF, which ultimately instructs proper nucleosome positioning along the fiber, perhaps through recruitment of ISWI-family chromatin remodelers^42,43^. When CTCF itself is no longer present, however, mature nucleosomes occupy CTCF motifs at steady-state, and upon replication, these fibers become hyperaccessible as they are repopulated by (parental or recycled) underwound nucleosomes and/or subnucleosomal particles. This model points to the incredibly fidelity and tight regulation of histone recycling and deposition coupled with replication fork movement, which together accommodate multiple pathways of chromatin fiber assembly post-replication, in a TF-dependent manner.

## DISCUSSION

We present RASAM, a long-read sequencing method that measures single-molecule nascent mammalian chromatin accessibility. Our pilot datasets provide the highest-resolution views to date of individual nucleosomes and TFs on nascent chromatin fibers at genome scale, including within ‘unmappable’ repetitive regions. Given the broad utility of nucleotide-analog-labeling, we anticipate application of RASAM to myriad topics, including the study of lagging- / leading-strand-coupled chromatin fiber assembly, how individual chromatin remodelers mediate this process, and how chromatin is remodeled by replication stress. Our study also raises the possibility of applying RASAM *in vitro*: we envision that reconstitution of replication on chromatin templates from genomic sequences will enable us to quantify how individual proteins contribute to the observed pattern at these loci post-replication *in vivo*^25^. Finally, incorporating genome-wide RASAM measurements into highly-resolved predictive models of genome activity^38,44^ and structure^45^ presents a promising avenue for future research for understanding epigenetic memory in cellular differentiation.

We note multiple areas to further develop our method. RASAM does not currently enrich for BrdU-containing molecules; instead, libraries are subjected to deep PacBio sequencing, with BrdU^(+)^ fraction dependent on labeling time. While this has attendant benefits (*e.g.* direct comparison of steady-state and nascent molecules from the same *in vivo* footprinting reaction), incorporating biochemical enrichment of BrdU^(+)^ dsDNA^5^ into the RASAM protocol will enable routine single-molecule study of the nascent genome at higher coverage. Computationally, we highlight several areas for improvement in our proof-of-concept pipeline, including base-pair and strand-specific resolution of BrdU, and multiplexed detection of other halogenated nucleotide analogs (*e.g.* CldU, IdU). These improvements will extend the biological scope of RASAM to measure replication fork speed and direction concurrently with single-molecule chromatin accessibility.

### Nascent mammalian genomes are hyperaccessible

Applying RASAM in both mESCs and human K562 cells, we observe a single-molecule chromatin state we term “nascent chromatin hyperaccessibility,” a transient state that is resolved over several hours of chromatin maturation to yield steady-state accessibility patterns. This discovery ties back to initial biochemical observations of chromatin at reconstituted and native replication forks, which demonstrated negatively-supercoiled chromatin hypersensitive to MNase digestion or hyperaccessible to acetyltransferases^3,46,47^. Our nondestructive, genome-wide, single-molecule measurements provide essential clarity into structure suggested by these assays: we demonstrate that the average nucleosome on nascent chromatin fibers is either underwrapped or subnucleosomal (*e.g.* a hexasome), and that nascent fibers are enriched for irregularly-spaced nucleosomes.

What processes drive and resolve nascent chromatin hyperaccessibility? At the length-scale of individual nucleosomes, there are multiple possible sources: RASAM may be capturing nucleosome core particle assembly intermediates consequent to activities of CAF-1^48^ and other histone chaperones; nascent nucleosomes and / or subnucleosomes may be more favorable substrates for ATP-dependent (*e.g.* INO80)^49^ or -independent (*e.g.* histone acetyltransferases, polycomb repressive complexes [PRC] 1 and 2)^47,50^ factors due to their organization (*i.e.* NRL and/or their irregularity); or, torsional strain induced by the replisome may allow nucleosomes to significantly underwrap without disassembly^51^. At the length-scale of individual fibers, we provide evidence that nucleosome positions on irregularly-spaced nascent arrays are subtly—but significantly—more constrained by dinucleotide content than steady-state arrays, implying that nascent chromatin assembly is initially more influenced by sequence-directed nucleosome positioning than those fibers that have undergone chromatin maturation. Combining RASAM with acute combinatorial perturbation of topoisomerases, ATP-dependent chromatin remodelers, and other replication-associated proteins will ultimately clarify pathways responsible for nascent chromatin hyperaccessibility.

### Understanding regulatory specificity in newly replicated chromatin

Our RASAM measurements paint nascent chromatin as a unique regulatory substrate. Nascent fibers are composed of immature nucleosomal species that remain hyperaccessible hours after replisome passage for various epigenomic domains that we analyzed. In contrast, we show that multiple TF motifs and TSSs on nascent chromatin are not more accessible than steady-state chromatin at these sites. Our data at CTCF and TSSs suggest distinct routes to re-establishing active *cis*-regulatory elements post-replication. At CTCF sites, factor occupancy and motif accessibility are reduced immediately after replication and return to steady-state levels after an hour; targeted degradation experiments demonstrate CTCF itself regulates this program, as factor depletion restores nascent chromatin hyperaccessibility. At TSSs, NFRs are re-established rapidly behind the fork; this implies that nucleosome assembly behind the replication fork is either inhibited at NFRs, or that chromatin remodeling activities immediately space and evict assembled nucleosomes post-replication. These data imply that the replication machinery and other associated protein complexes (*e.g.* histone chaperones responsible for depositing nucleosomes) are uniquely regulated at specific, replicating *cis*-regulatory loci.

Prior studies have demonstrated that ATP-dependent chromatin remodelers^6^ and RNA Pol II transcription^9^ dictate nascent chromatin accessibility at active regulatory regions. Our results add to this list by showing that CTCF directly regulates nascent chromatin structure at its binding motifs. How this might occur remains an exciting mystery. TF binding on pre-replicated chromatin could reinforce high local-concentrations of necessary co-factors (*e.g.* ISWI remodelers for CTCF)^42,43^ or directly regulate histone deposition behind the fork, thus programming a post-replicative pattern of chromatin maturation; alternatively, the bound TF may directly signal to the replisome itself (*e.g.* through DNA mechanics), through steric occlusion of a traveling fork, or via other specific protein-protein interactions. Future work dissecting specific CTCF-associated regulatory proteins will provide insight into this process.

The non-destructive nature offered by RASAM reconciles many seemingly contradictory aspects of chromatin maturation behind the replication fork. First, we show that the majority of newly-replicated chromatin fibers harbor well-spaced nucleosome core particles, but that nascent fibers are statistically-enriched for fibers lacking regularity; our data are thus consistent with classical results describing fast chromatin reassembly post-replication^52^, and a model in which primary DNA sequence transiently influences nucleosome positioning behind the fork. Second, we find that the nascent genome is globally hyperaccessible; though this contradicts one conclusion from population-averaged repli-ATAC-seq (possibly owed to differences between accessibility measurements using Tn5 insertion versus EcoGII methylation), our single-molecule measurements of rapidly reformed NFRs support the finding that active transcription instructs chromatin maturation^9^. Third, though we find reduced CTCF occupancy on individual nascent fibers (consistent with competition between nucleosomes and TFs on nascent chromatin, and lack of Tn5 hyperaccessibility at nascent CTCF motifs)^6,9^, we also observe individual newly-replicated chromatin fibers with CTCF footprints and phased nucleosomes, though we note that we cannot formally assess potential ‘bookmarking’ functions for CTCF and other TFs behind the replication fork^13^ using RASAM.

Finally, our results question the notion that primary template accessibility alone can specify genome regulation. Eukaryotic gene regulation is predicated on the accurate sorting of sequence-dependent and -independent (co)activators to specific *cis*-regulatory elements, and away from cryptic binding sites in heterochromatin. Our observation of a hyperaccessible genome upon DNA replication would suggest that multiple regulators are necessary to preserve *cis*-regulatory landscapes across divisions with high fidelity. Considering this, and integrating our results with the literature, we propose a ‘tuning’ model for chromatin maturation behind the replication fork (**Figure 7**). In this model, high-fidelity maintenance of regulatory states is enforced by the hierarchical activities of specific regulatory factors (*e.g.* histone chaperones, ATP-dependent chromatin remodeling, topoisomerase activity, transcription) as a function of ‘maturation time’ post-replication. By specifying maturation dynamics for various classes of binding sites (*e.g.* promoters, CTCF sites, enhancers), distinct groups of regulators working in a stepwise manner can support accurate maintenance of regulatory programs, while minimizing cryptic factor binding and heterochromatin erosion.

**Figure 7:**
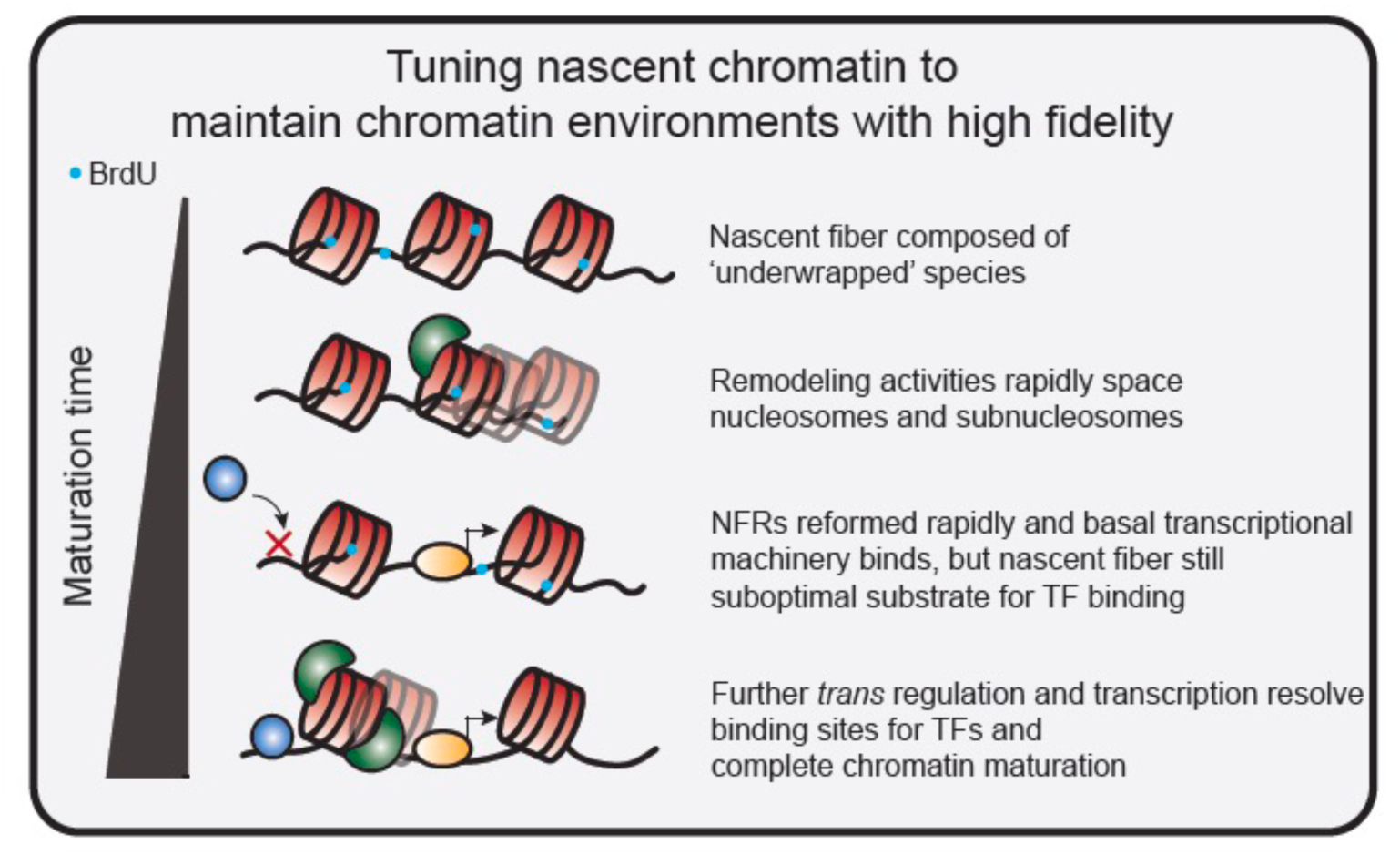
A proposed model for nascent chromatin fiber maturation, integrating results of this study and others. We propose a ‘tuning’ model for nascent chromatin maturation, in which specific regulatory factors dictate a hierarchy of events underlying faithful chromatin maturation across cell cycles. In this model, remodeling activities rapidly space nucleosomes and reset the nucleosome free region of active promoters. While transcription is able to restart, steady-state levels of binding for other factors is diminished as the nascent fiber still represents a suboptimal binding substrate. Finally, additional *trans* regulators and the act of transcription, together resolve the chromatin landscape to accommodate steady-state TF binding and complete chromatin maturation.

## SUPPLEMENTARY FIGURES

**Supplementary Figure 1:**
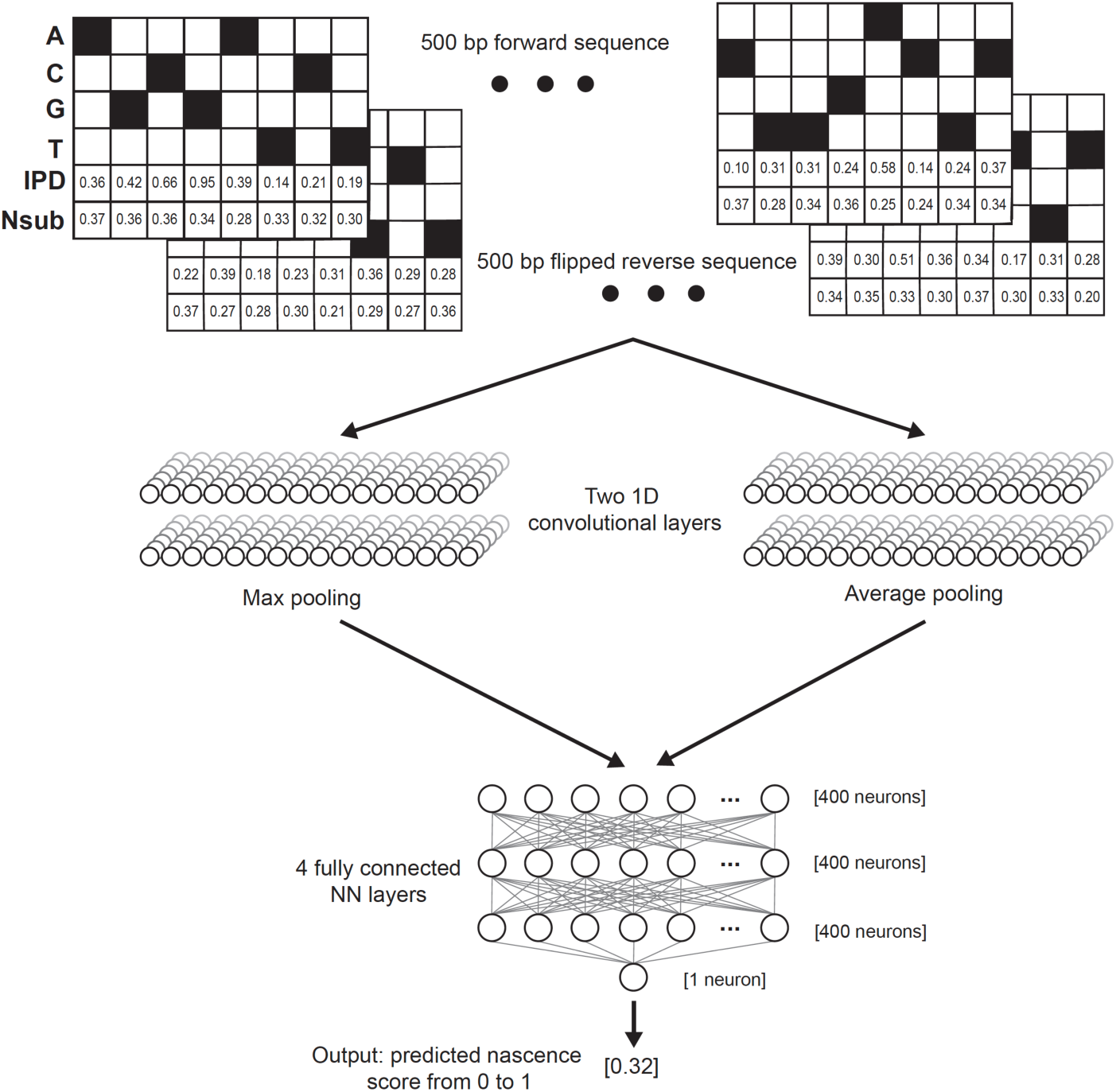
Schematic of the convolutional neural network (CNN) used to predict BrdU-containing molecules in this study. We engineered a CNN that takes a 500 bp ‘tile’ of one-hot encoded sequence (forward and reverse), along with the interpulse duration (IPD) for each base as reported by the PacBio Sequel II sequencer and the number of passes (Nsub), and predicts a scalar ‘nascence’ score for the input tile. The model is comprised of two parallel 1D convolutional layers that use max and average pooling, respectively, the outputs of which are fed into 4 fully connected neural network layers, each with 400 neurons.

**Supplementary Figure 2:**
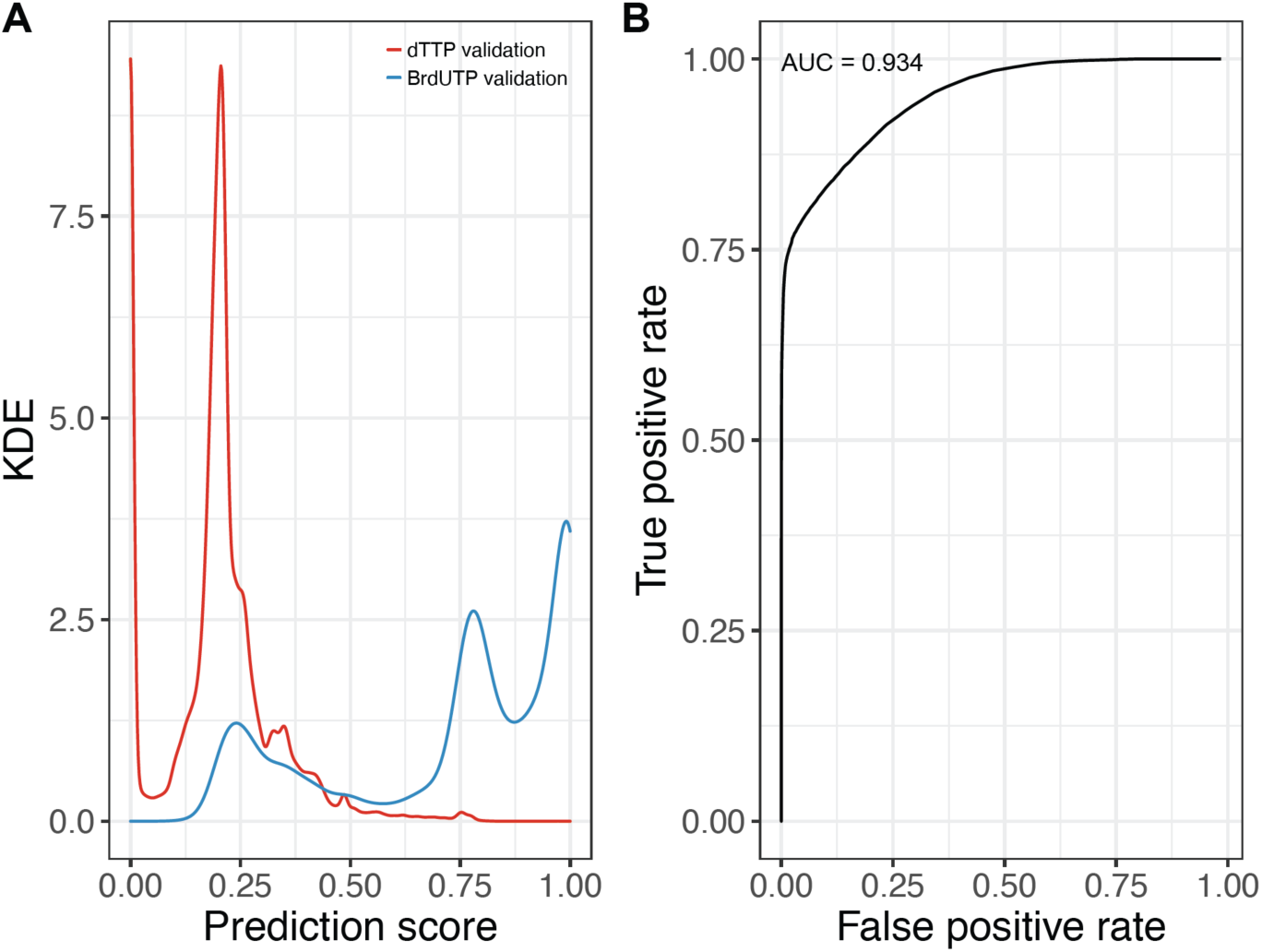
A convolutional neural network (CNN) enables prediction of BrdUTP-incorporation. **A.)** BrdU prediction score distribution for unlabeled negative control (red) and labeled positive control (mixture of PCR and *in vivo* labeled BrdUTP-containing molecules; blue) molecules, all of which were held-out from model training. **B.)** ROC curve of true positive rate as a function of false positive rate for CNN classifications. ROC curve demonstrates an AUC of 0.934.

**Supplementary Figure 3:**
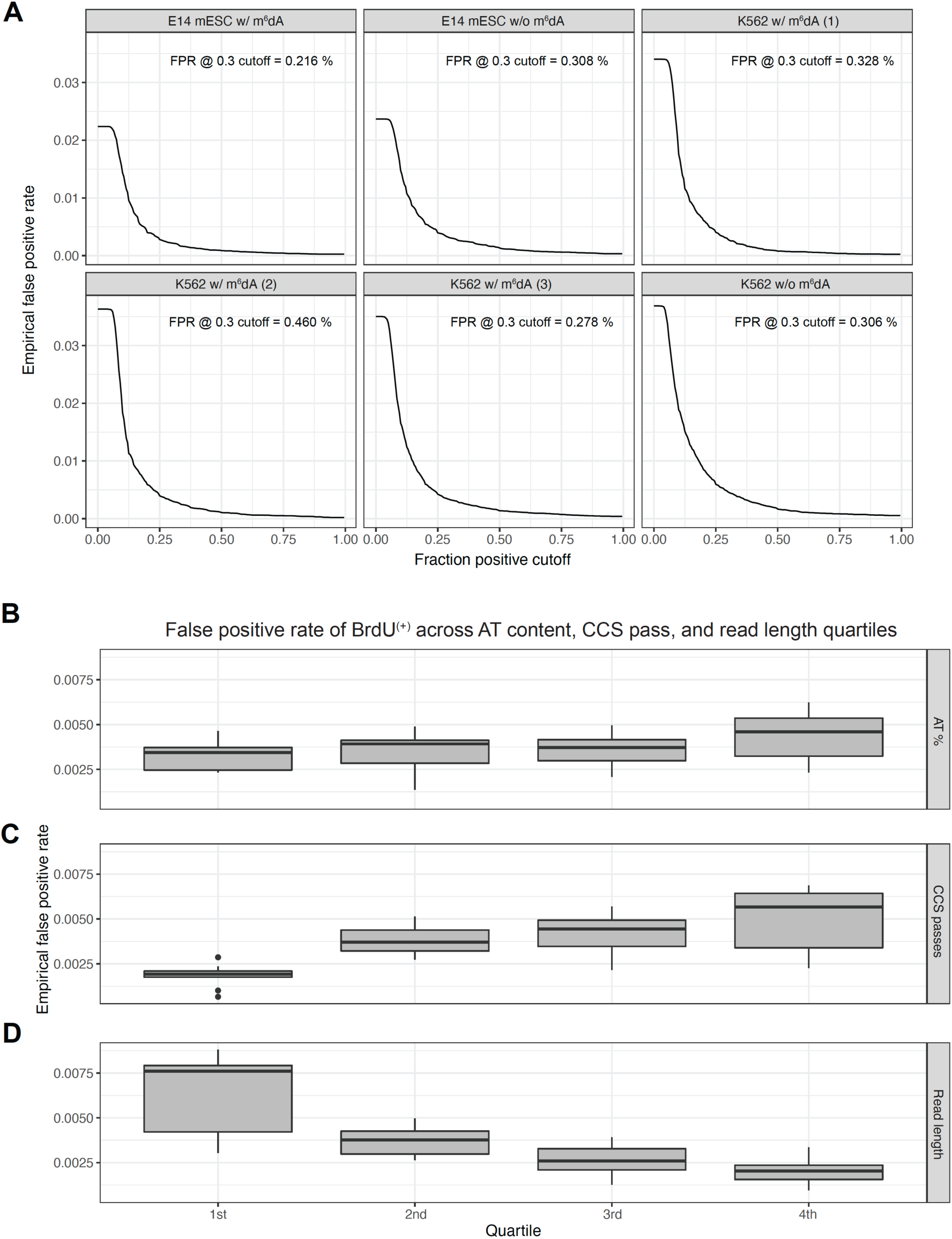
Defining an empirical false positive rate for RASAM BrdU(+) classification. **A.)** Using unlabeled negative control samples from K562 and E14, we defined a cutoff of a minimum fraction of positively-classified tiles (*i.e.* ‘fraction positive cutoff’), which would reproducibly predict a tolerably low BrdU(+) fraction in unlabeled samples. Here, we plot the empirical false positive rate as a function of the fraction positive cutoff for 2 E14 mESC samples and 4 K562 samples, both with and without m^6^dA footprinting. **B-D.)** False positive rate for samples for molecules stratified by AT-content (**B**), number of PacBio CCS passes (**C**), and read length (**D**). False positive rate is not influenced substantially by AT content of sequenced molecules, but is impacted by CCS passes and read length, which are inter-related. This is consistent with more uncertainty in classification for molecules with few CCS passes (including the longest sequenced molecules in datasets).

**Supplementary Figure 4:**
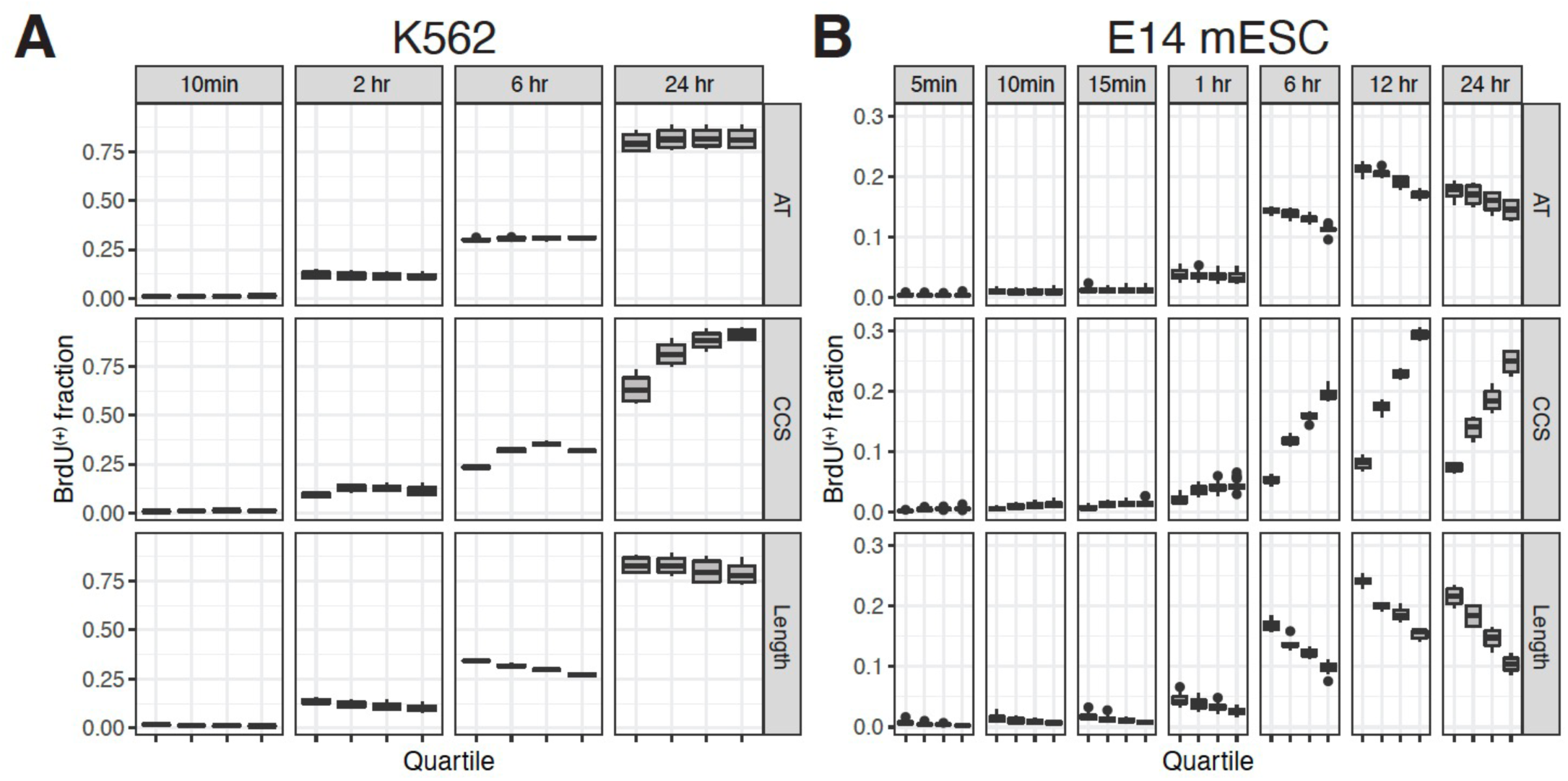
BrdU^(+)^ fraction is influenced primarily by CCS passes and read length. BrdU^(+)^ fraction for labeling experiments in K562 (**A**) and mESC (**B**) experiments, stratified by quartiles of AT-content (top), CCS passes (middle), and read length (bottom). We observe length and CCS pass-dependent drops in BrdU^(+)^ fraction consistent with increased classification uncertainty in the longest sequenced molecules (which also generally have the fewest CCS passes).

**Supplementary Figure 5:**
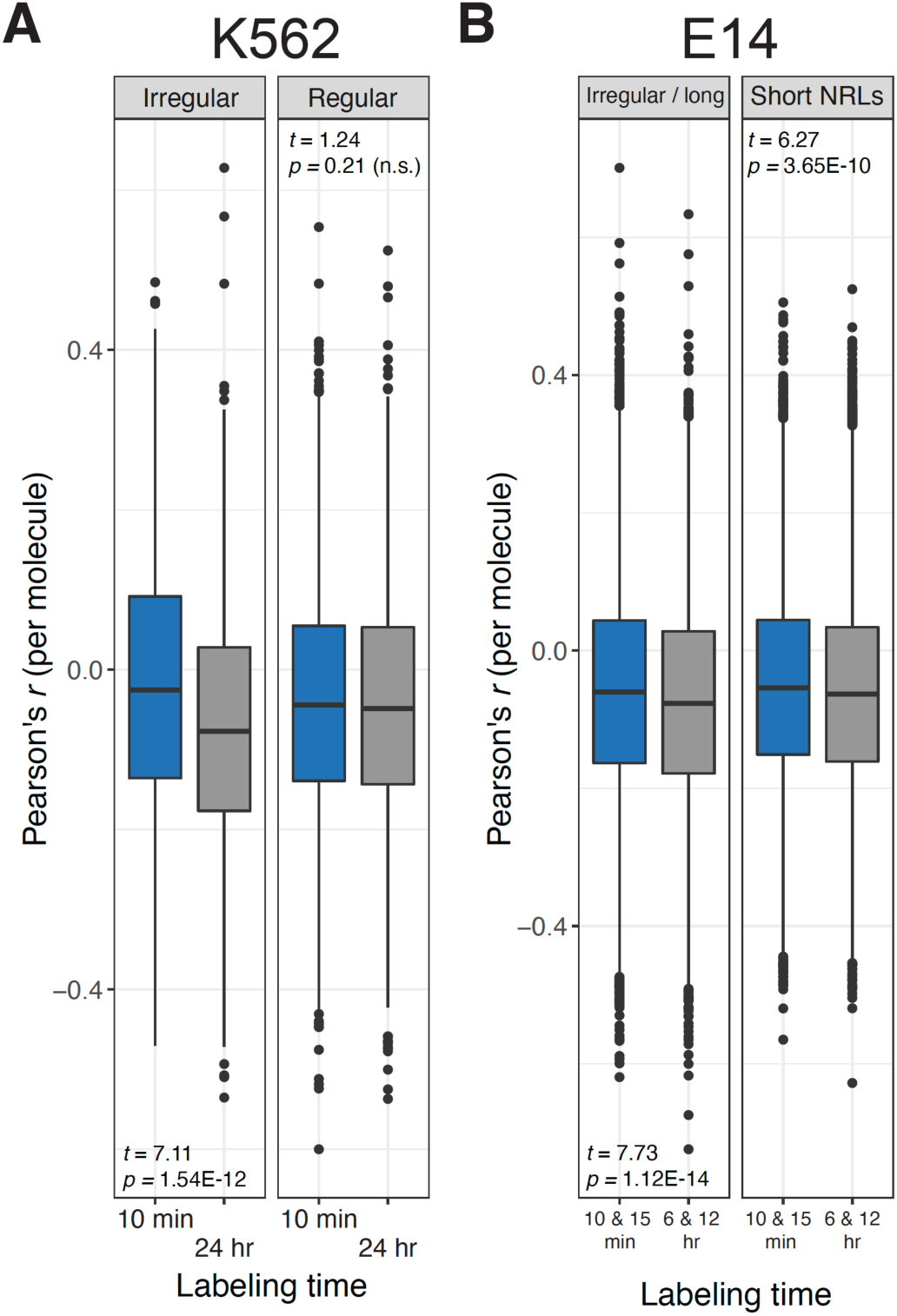
Irregularly-spaced nascent fibers are significantly more correlated with primary sequence compared to irregularly-spaced long-time point labeled fibers. **A.)** Boxplot representation of the per-molecule Pearson’s correlation (*r*) between RASAM accessibility signal and rolling AT-dinucleotide content for irregularly-spaced BrdU^(+)^ fibers after labeling for 10 min (blue; ‘nascent’) and 24 hr (grey; ‘steady-state’). Single-molecule correlations are significantly higher (student’s *t*-test *p* = 1.54E-12) in irregularly-spaced nascent fibers, compared to irregularly-spaced steady-state fibers. For comparison, there is no significant difference in correlations for regularly spaced fibers between these two time points (student’s *t*-test *p* = 0.21). **B.)** As in (**A**), but in E14 mESCs, for irregularly-spaced and long NRL fibers (left) and short NRL regularly-spaced fibers (right). In both cases, we observe subtle but significant increases in per-molecule Pearson’s correlation between nascent (10 min / 15 min labeling) versus long-time point labeling (6 hr / 12 hr) fibers (left student’s *t*-test *p* = 1.12E-14; right student’s *t*-test *p* = 3.65E-10), indicating increased influence of sequence on single-molecule accessibility in nascent fibers.

**Supplementary Figure 6:**
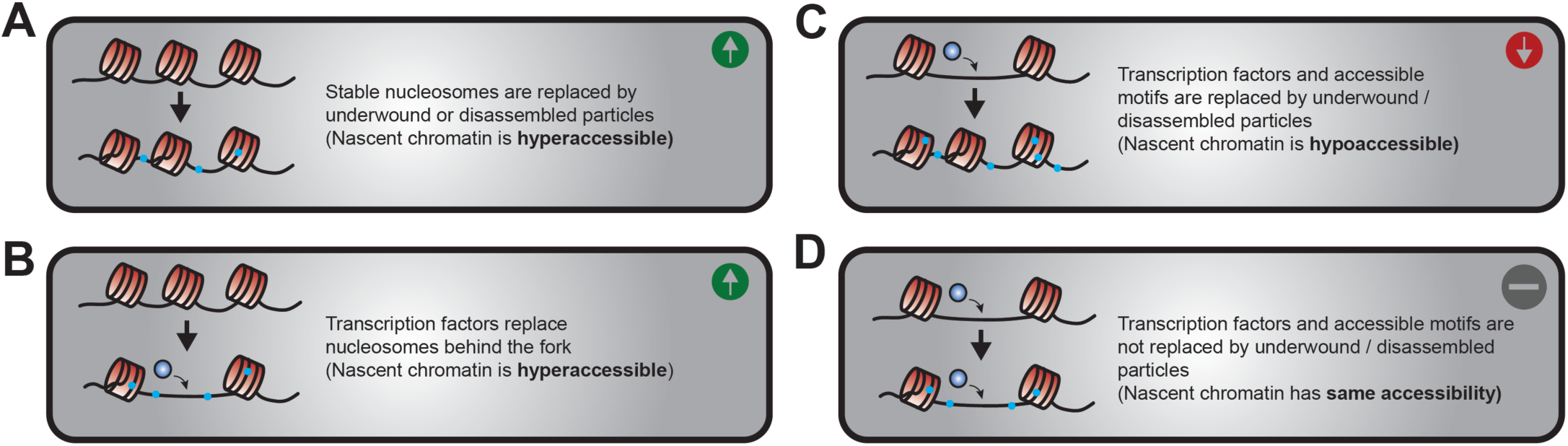
Four possible models for how individual chromatin fibers may assemble post-replication, accounting for nascent chromatin hyperaccessibility. **A.)** Genome-wide, nascent chromatin hyperaccessibility can be simply attributed to the replacement of stable nucleosomes with underwound nucleosomal species behind the fork. **B.)** At active and cryptic *cis-*regulatory motifs, nascent chromatin hyperaccessibility could occur if mature nucleosomes are not recycled or deposited post-replication, allowing TFs to bind cognate motifs. **C.)** Nascent chromatin can display hypoaccessibility if TFs and nascent nucleosomes compete for the same binding sites behind the fork. **D.)** Nascent and steady-state chromatin fibers could display similar single-molecule accessibility if nucleosome-free regions are preserved (actively or passively) immediately post replication.

**Supplementary Figure 7:**
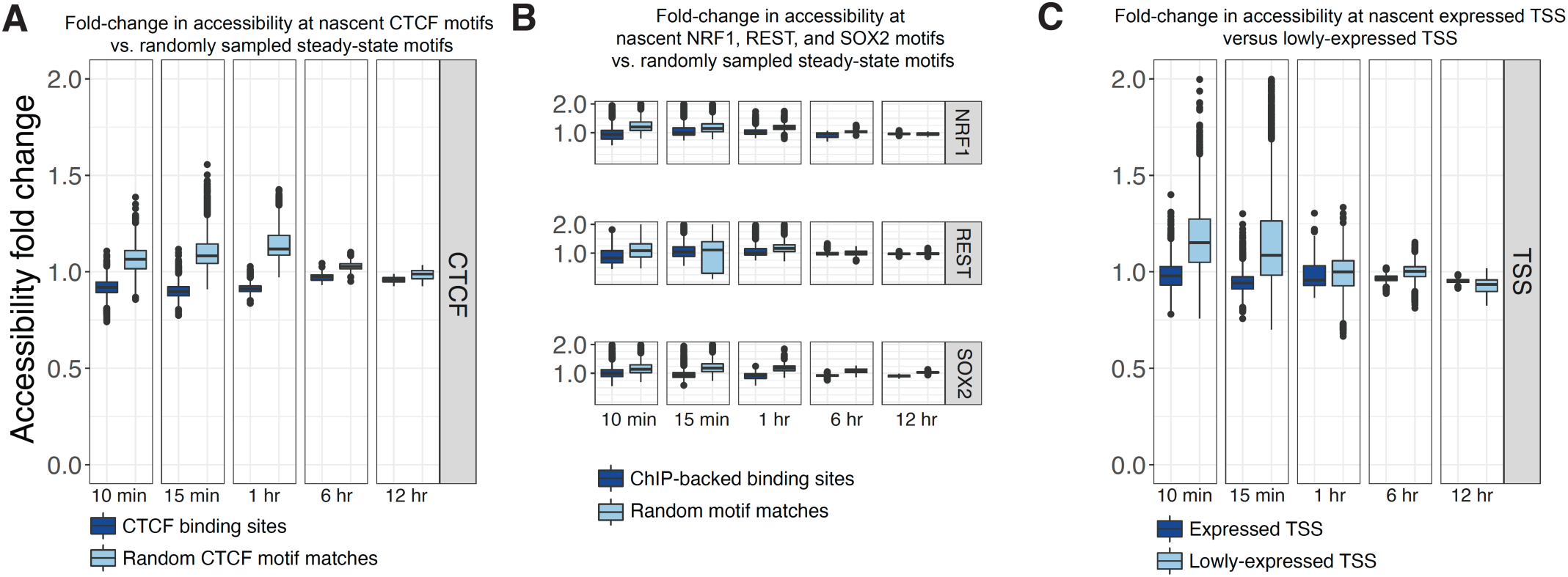
Nascent chromatin hyperaccessibility is constrained at active TF binding sites. **A.)** Box plots of fold change accessibility estimates between BrdU^(+)^ and unlabeled molecules at CTCF motifs for ChIP-backed binding sites (dark blue) and random CTCF motif matches (light blue), stratified by labeling time. Box plot distributions represent the median and interquartile range for accessibility fold changes calculated from the average accessibility of BrdU^(+)^ molecules divided by *n* = 1,000 randomly sampled, equivalent numbers of unlabeled molecules per biological replicate (*n* = 6,000 values per boxplot). **B.)** As in (**A**) but for NRF1 (top), REST (middle), and SOX2 (bottom). **C.)** As in (**A-B**) but comparing the TSSs for the top 80% expressed genes (dark blue) against the bottom 20% of expressed genes (light blue; unexpressed genes omitted in this analysis) in mESC.

**Supplementary Figure 8:**
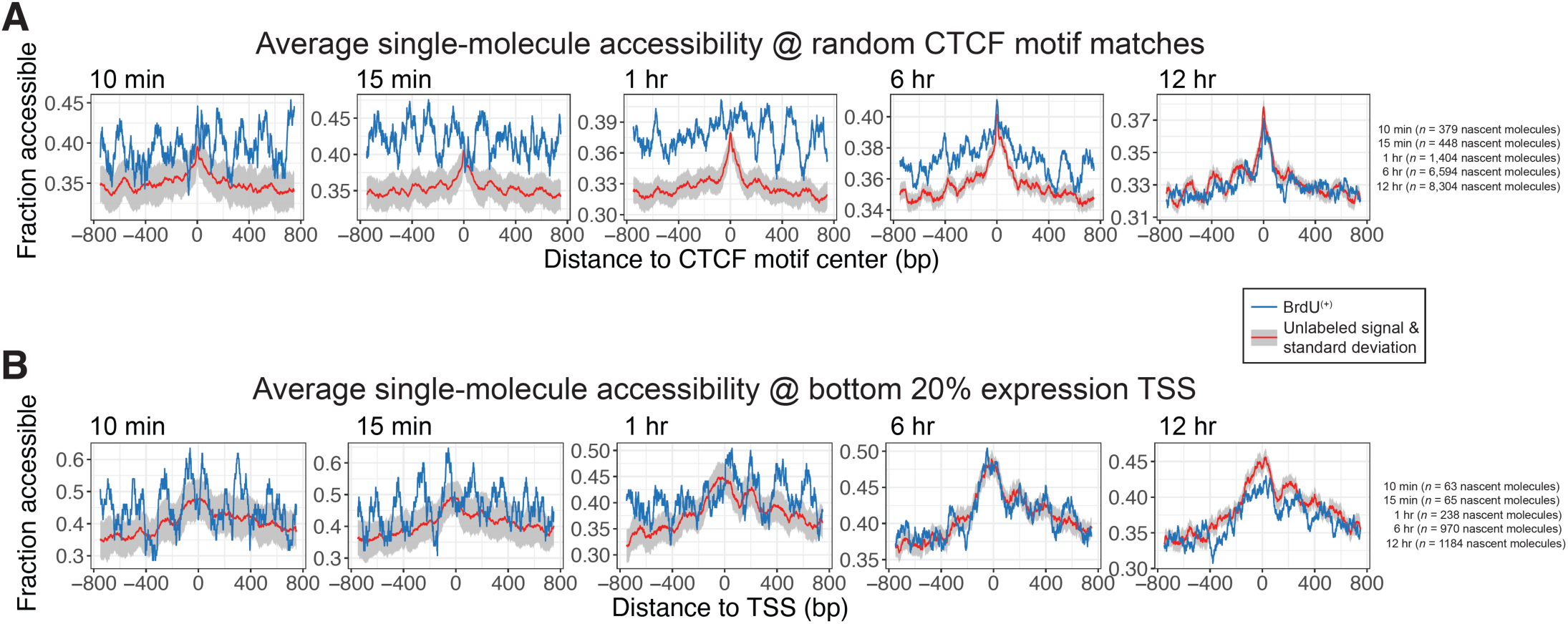
Average single-molecule accessibility at negative control sites for CTCF and TSSs for the lowest 20% expressed genes in mESCs. **A.)** Line plot visualization of average single-molecule accessibility of BrdU^(+)^ (blue) and unlabeled (red) fibers centered at random matches to the CTCF motif in the mm10 mouse genome assembly, stratified by labeling time. Grey ribbons around unlabeled signal represent the standard deviation of mean accessibility calculated from *n* = 100 random samples of an equivalent number of unlabeled molecules, compared to BrdU^(+)^ molecules. **Inset:** tabulated BrdU^(+)^ fiber counts for each labeling time point. **B.)** As in (**A**), but for fibers overlapping with a TSS for the bottom 20% lowest expressed genes in mESC.

**Supplementary Figure 9.**
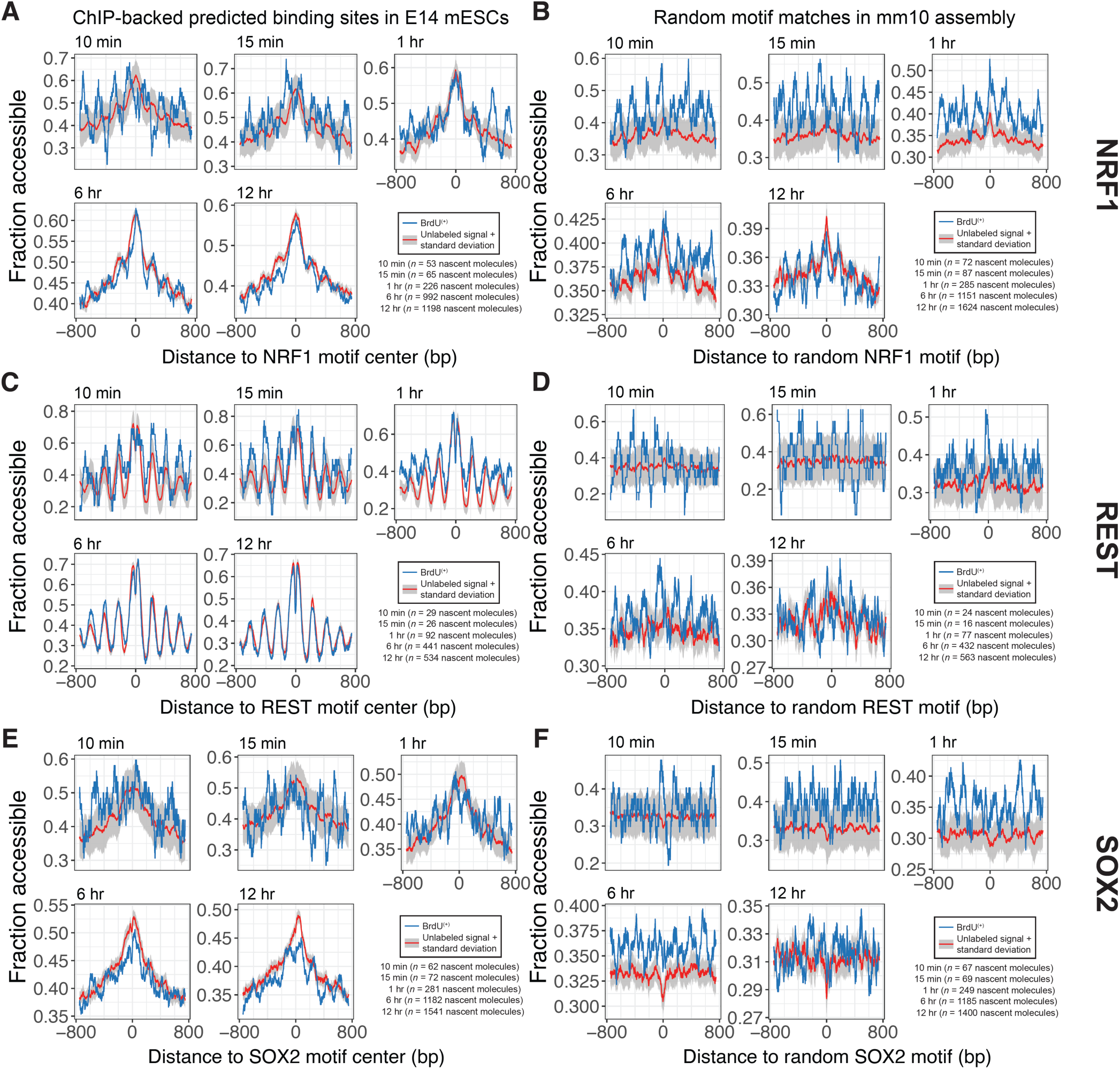
Average single-molecule accessibility at assorted TF binding sites in the mESC epigenome. **A.-B.)** Line plot visualization of average single-molecule accessibility of BrdU^(+)^ (blue) and unlabeled (red) fibers centered at ChIP-backed NRF1 binding sites (**A**) and random Nrf1 motif matches (**B**), stratified by labeling time. Grey ribbons around unlabeled signal represent the standard deviation of mean accessibility calculated from *n* = 100 random samples of an equivalent number of unlabeled molecules, compared to BrdU^(+)^ molecules. **Inset:** tabulated BrdU^(+)^ fiber counts for each labeling time point. **C-D.)** As in (**A-B**) but for REST. **E-F.)** As in (**A-B**) but for SOX2.

**Supplementary Figure 10.**
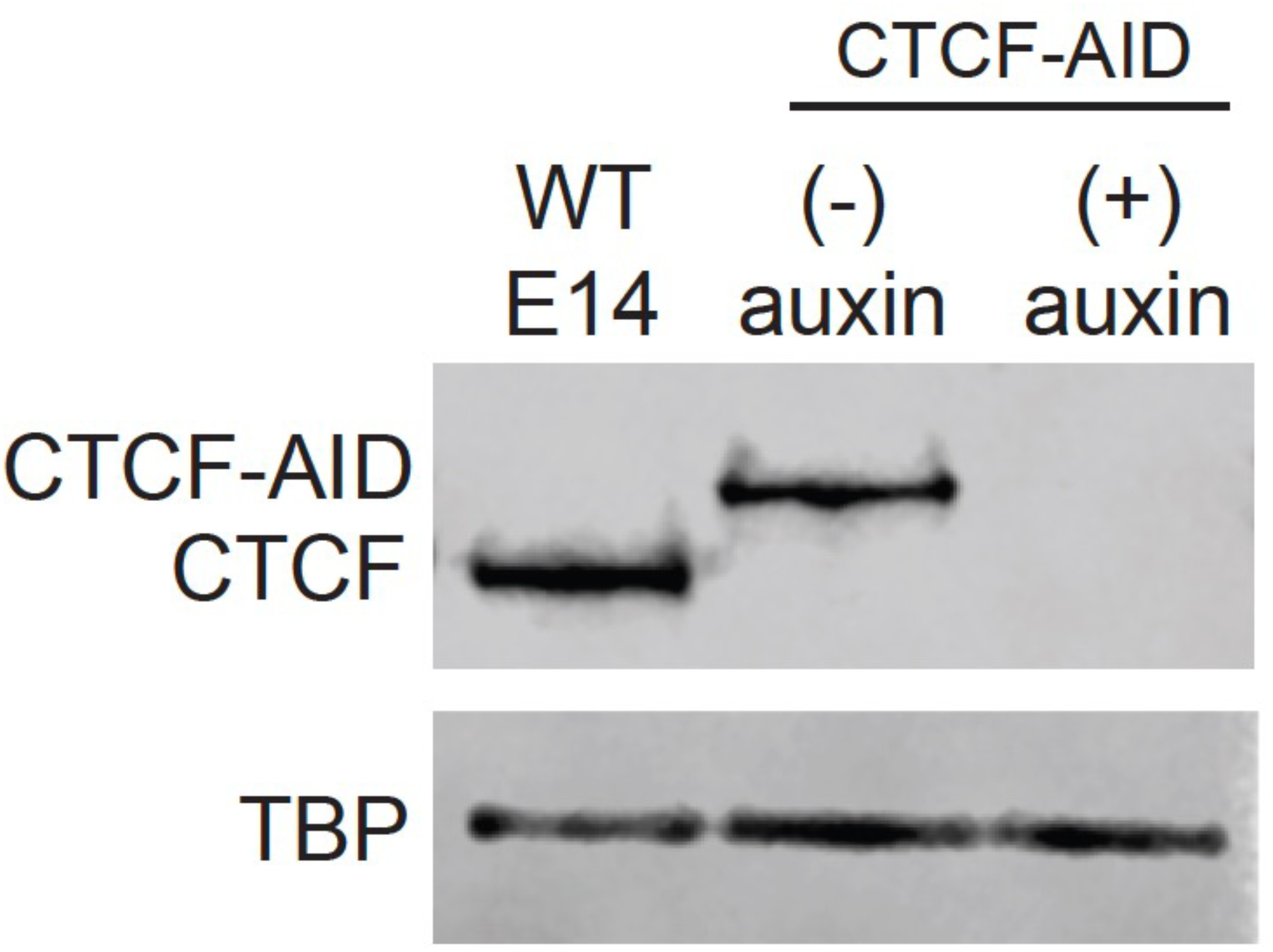
Western blot validation of CTCF degradation in CTCF-AID E14 mESCs. Lane 1: positive control of untagged, wildtype E14 mESCs; Lane 2: Untreated CTCF-AID E14 mESCs; Lane 3: Auxin-treated (6 hr) CTCF-AID E14 mESCs. TBP used as loading control for all samples.

**Supplementary Figure 11:**
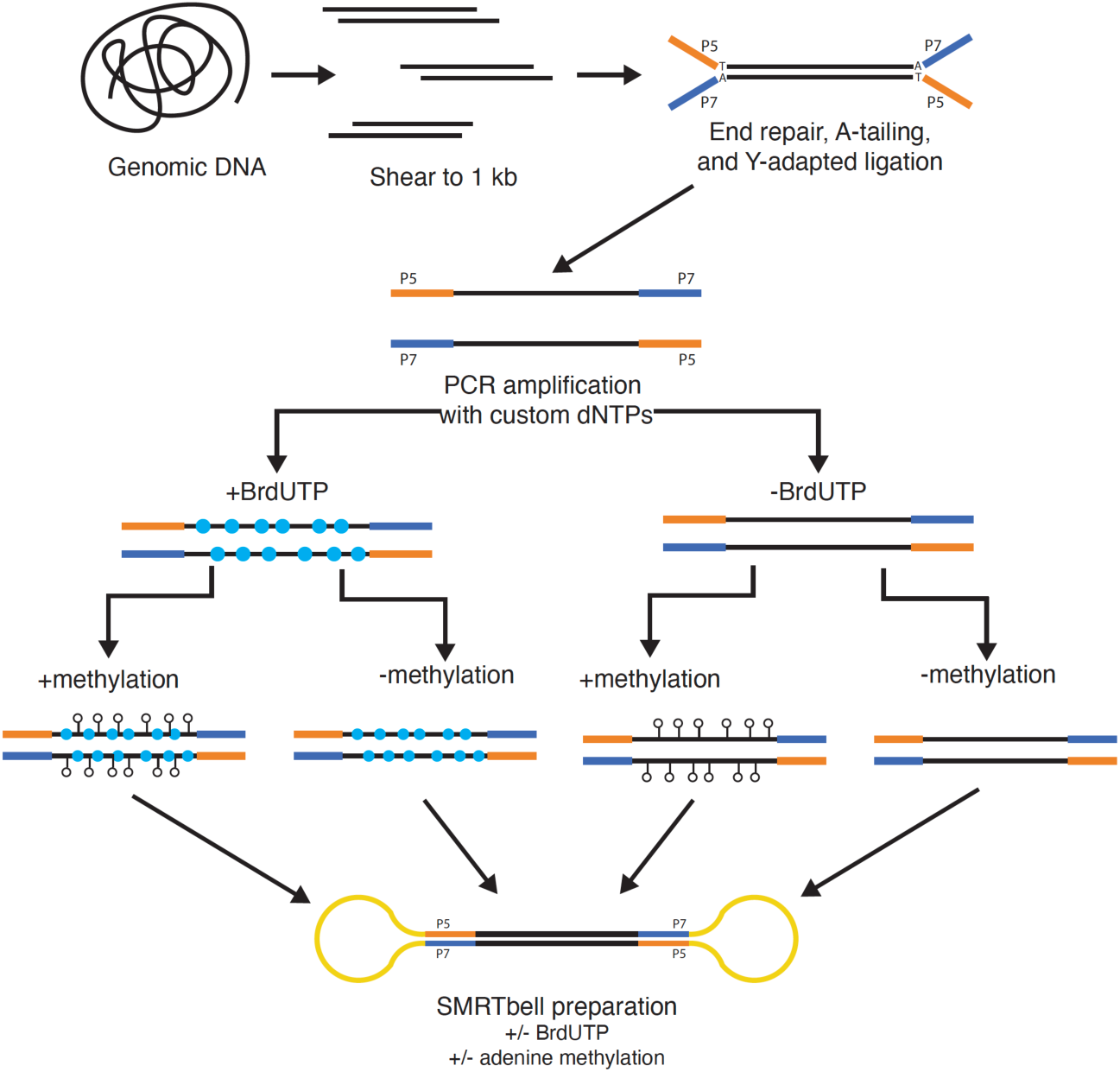
Strategy for generating PCR amplicons to train a BrdU-prediction model. We devised a PCR-based strategy employing Y-adaptor ligated mouse and human genomic DNA as a template to generate genome-derived amplicons with defined nucleotide content. Using uracil-tolerant polymerases, we generated templates with 100% BrdUTP, a mixture of BrdUTP and dTTP, and 100% dTTP, all of which were used along with genomic DNA samples to train the convolutional neural network.

**Supplementary Figure 12.**
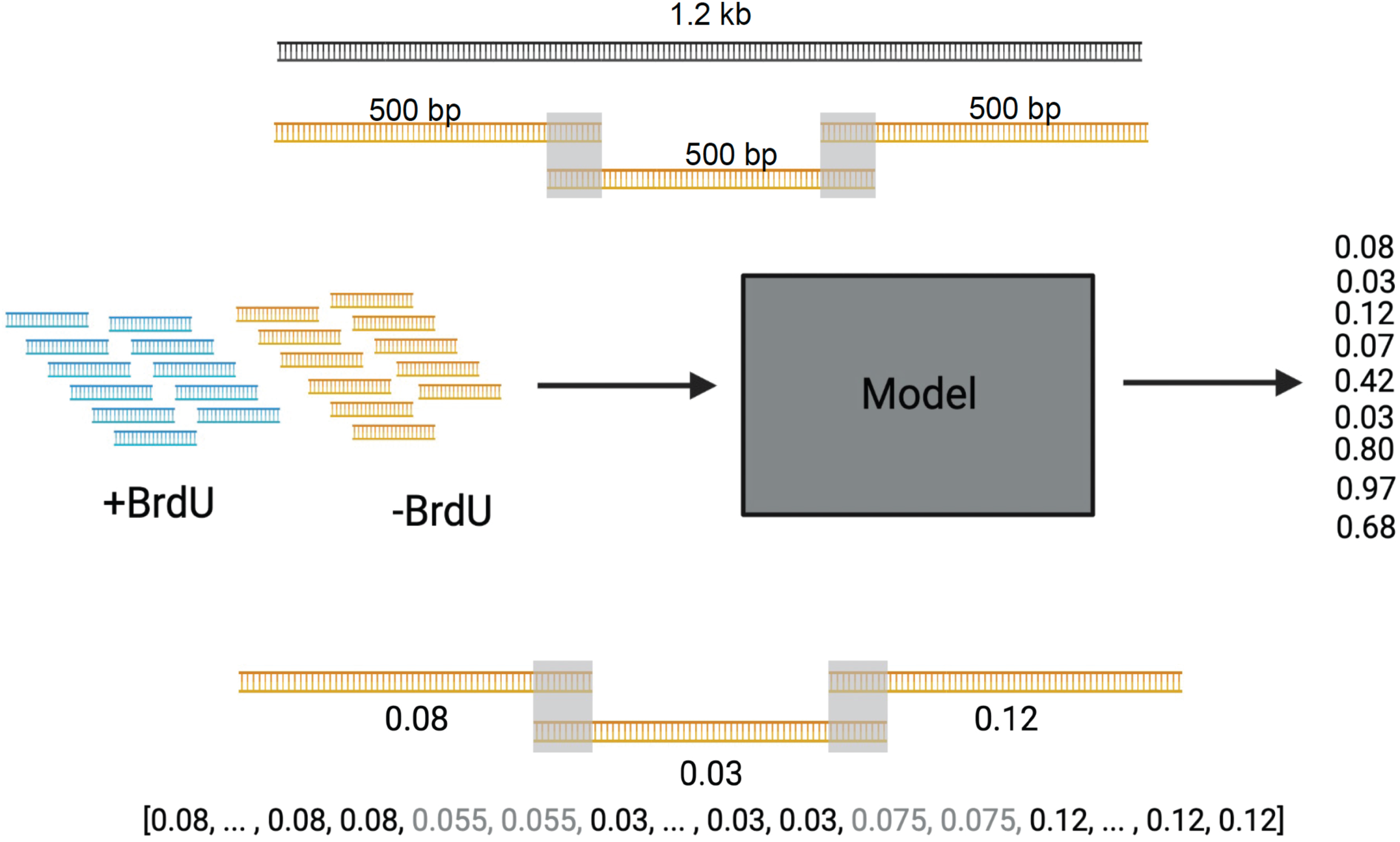
Schematic of how 500 bp BrdU^(+)^ classifications were lifted over to full-length molecules. After classifying 500 bp sequences, classifications were ‘stitched’ back to parent molecules, and regions of overlap between 500 bp classifications were averaged, as shown in the cartoon above.

## METHODS

### Cell Lines and Cell Culture

K562 cells (ATCC) were grown for at least three days in standard media containing RPMI 1640 + L-Glutamine (Hyclone) supplemented with 10% Fetal Bovine Serum (Gibco) and 1% Penicillin-Streptomycin (Gibco). mESC E14 cells were grown on 0.2% gelatin and maintained in standard media containing DMEM + Glutamax (ThermoFisher 10566-016), 15% Fetal Bovine Serum (ThermoFisher SH30071.03), 1X non-essential amino-acids (ThermoFisher 11140-50), 1 mM Sodium pyruvate (ThermoFisher 11360-070), 0.128mM 2-Mercaptoethanol (BioRad 1610710XTU), and 1X Leukemia Inhibitory Factor (purified and gifted by Barbara Panning Lab at UCSF). MESCs were passaged every 1-2 days using Tryple Express Enzyme (1X) (12605010). Media was changed daily when cells were not passaged. All cell lines were regularly tested negative for mycoplasma contamination.

### Western Blotting

Degron line cells were treated with 500 uM Indole-3-acetic acid sodium salt (Sigma I5148) for 6 hours prior to collection. 10-20 million cells were used to prepare whole cell lysates. Cells were dissociated using Tryple Express Enzyme (1X) (12605010), resuspended in cell media, spun down (300xg for 5 min), washed in 1X PBS and spun again. Cell pellets were resuspended in 100 uL cold RIPA buffer (150 mM NaCl, 1% NP-40, 0.5% SDS, 50mM Tris-Cl, pH 8.0, and 1X Protease Inhibitor (Roche 4693132001)) per 1 million cells. The cell suspension was agitated at 4°C for 30 min, then sonicated in Covaris microTUBE AFA Fiber tubes (6×16mm). Sonicated lysate was centrifuged at 12,000 RPM for 20 mins at 4°C and protein concentration was measured using Pierce BCA Protein Quantification. 25 ug of whole cell lysate was loaded per lane. Samples were mixed with NuPage LDS buffer (ThermoFisher NP0007) and 5% b-mercaptoethanol, then heated at 100°C for 5 minutes and loaded onto a 4-12% polyacrylamide gel (ThermoFisher NP0322). Transfer onto a PVDF membrane was performed in 1X NuPage Transfer Buffer (ThermoFisher NP00061) with 0.1% SDS for 60 minutes at 30V and the membrane was subsequently blocked in 5% Bovine Serum Albumin in 1x PBS with 0.1% Tween-20 (PBST) overnight at 4°C. Primary antibody staining was done at manufacturer’s recommended dilution in 1x PBST for 1 hour at room temperature followed by 3 rinses at room temperature with 1x PBST. The secondary antibody stain was done at room temperature for 1 hour at a 1:10,000 dilution in 1x followed by 3 rinses with 1x PBST. The membrane was incubated in WesternSure PREMIUM Chemiluminescent Substrate (LiCor) for 5 minutes before imaging on a Li-Cor imaging system.

### Nucleotide Analog Incorporation in vitro

Genomic DNA from mESC E14 or K562 cells was MNase digested or sheared to ∼1 kb and then end repaired, A-tailed, and adapter ligated to then be used as input for PCR. PCR was performed with Q5U HotStart High-Fidelity polymerase (New England Biolabs, M0515) as usual except a custom dNTP mix was used that contained dATP, dCTP, dGTP, and either 100% dTTP, 100% BrdUTP, or 50% dTTP and 50% BrdUTP. The resulting DNA fragments were then either methylated using non-specific adenine EcoGII methyltransferase (New England Biolabs, high concentration stock 2.5e4 U/mL) or left unmethylated. All samples were then used as input for PacBio library prep using the SMRTbell Express Template Prep Kit 2.0.

### RASAM

#### Nucleotide labeling

Standard media was added to mESC E14 or K562 cells with a final concentration of 10 uM BrdU (abcam ab142567). Cells were incubated in BrdU-containing media for between five minutes and 24 hours. After labeling, mESC E14 plates were rinsed three times with 1x PBS and collected using Tryple Express Enzyme (1X) (12605010). K562s were collected and spun down (300xg, 5 minutes) immediately after labeling.

#### Nuclei isolation

Briefly, all nuclei were collected by centrifugation (300xg, 5 min), washed in ice cold 1X PBS, spun again (300xg, 5 min), and resuspended in 1 mL Nuclear Isolation Buffer (20mM HEPES, 10mM KCl, 1mM MgCl2, 0.1% Triton X-100, 20% Glycerol, and 1X Protease Inhibitor (Roche 4693132001)) per 10 million cells by gently pipetting 5x with a wide-bore pipette tip. The suspension was incubated on ice for 5 minutes, and nuclei were pelleted (600xg, 4°C, 5 min), washed with Buffer M (15mM Tris-HCl pH 8.0, 15 mM NaCl, 60mM KCl, 0.5mM Spermidine), and spun once again (600xg, 4°C, 5 min).

#### EcoGII footprinting

Nuclei were resuspended in 200 uL of Methylation Reaction Buffer (Buffer M containing 1mM S-adenosylmethionine (SAM)) (New England BioLabs B9003S). 10uL high-concentration EcoGII was added per 1e6 nuclei and the nuclei suspension was incubated at 37°C for 30 minutes. An additional 1 uL of 32 mM SAM was supplemented after 15 minutes. Any unmethylated controls were treated similarly with the addition of SAM, but EcoGII was excluded from those reactions.

#### DNA purification

To the methylation reaction, 2.65 uL 10% SDS and 2.65 uL of 20 mg/mL Proteinase K (Thermo Scientific AM2548) were added and incubated at 65°C for at least 2 hours and up to overnight. To extract the DNA, an equal volume of Phenol-Chloroform was added and mixed vigorously by shaking. The samples were then spun at max speed for 2 min and the aqueous portion was removed. To the aqueous extraction, 0.1x volumes of 3M NaOAc, 1 uL GlycoBlue, and 3x volumes of cold 100% EtOH were added, mixed by inversion and incubated overnight at −20 °C or for 2 hours at −80°C. Samples were then spun at max speed for 30 minutes at 4 °C, washed with 500 uL 70% EtOH, spun again at max speed for 2 minutes at 4 °C. The resulting pellet was air dried and resuspended in 40 uL EB. Samples concentrations were measured via Qubit High Sensitivity DNA Assay.

#### Preparation of PacBio SMRTbell Libraries

Purified DNA from mESCs and K562s was sheared using a Covaris g-tube (520079) in a 5424 rotor at 7,000 RPM for 6 passes for a target size between 6,000 and 8,000 bp. Seared DNA was used as input for the PacBio HiFi SMRTbell Library Preparation protocol using the SMRTbell Express Template Prep Kit 2.0. Briefly, samples underwent removal of single stranded overhangs, DNA damage repair, end-repair, A-tailing, barcoded SMRTbell adapter ligation, and exonuclease cleanup according to manufacturer’s protocol followed by a 1X AMPure PB bead cleanup. Final sample concentration was measured via Qubit High Sensitivity DNA Assay and library size was measured on an Agilent Bioanalyzer DNA chip. Libraries were sequenced on PacBio Sequel II 8M SMRTcells.

### E14 CTCF-AID experiments

E14 CTCF-AID mESCs ^41^ were cultured identically to wildtype E14 mESCs. To deplete CTCF-AID before nucleotide analog labeling, cells were incubated in 0.5 mM auxin (IAA Sigma, I5148) for 6 hours prior to any BrdU treatment. Wild-type E14 mESCs and CTCF-AID mESCs were gifted from Elphege Nora Laboratory at UCSF.

### Data Processing for single-molecule chromatin accessibility

Sequencing reads were processed using software from Pacific Biosciences and custom scripts available at https://github.com/RamaniLab/SAMOSA-ChAAT, as previously described ^28^. Scripts resulted in three primary files per sample: an alignment of CCS reads to the reference genome (mm10), an accessibility prediction per CCS molecule (Viterbi path of HMM component of model), and a list of footprints, molecule locations, and footprint sizes, all per molecule. All of these files were used in conjunction for downstream analyses.

### Training a model to predict BrdU^(+)^ from PacBio kinetics

Input for the convolutional neural network (CNN) was generated by dividing each molecule into discrete 500 bp segments that contained the one-hot encoded sequence, IPD, and the number of subreads for both the forward and reverse strands. These 500 bp chunks were each fed into two arms with two 1-dimensional convolutional layers with 200 units, relu activation, and he uniform initialization: one with max pooling and one with average pooling. The output from those arms were combined and used as input for a fully connected neural network consisting of 4 layers. The first three layers were comprised of 400 units with relu activation, he uniform activation, and a 0.5 dropout. The final layer contained 1 unit with sigmoid activation. The output from the final layer was a value between 0 and 1 for the probability of the sequence to contain BrdU. Training data included negative control *in vivo* samples and *in vivo* samples labeled for 24 hours in mESCs and K562s, 100% BrdUTP PCR amplicons with and without adenine methylation (**Supplemental Figure 11**), and methylated gDNA from mESCs and K562s. 588,000 molecules were used for training and 252,000 were used for validation. The model was trained using Adam optimizer for 15 epochs with a batch size of 32.

### Classifying BrdU^(+)^ from 500 bp tiles

To go from BrdU^(+)^ classification of 500 bp regions to BrdU^(+)^ classification of entire molecules, we adopted a smoothing strategy (**Supplementary Figure 12**), wherein molecules were separated into overlapping 500 bp chunks, classified, and then reassembled, with overlapping nucleotides scoring as the mean of each classification. Whole molecules were then further classified as BrdU^(+)^ using two cutoffs: first, we thresholded BrdU^(+)^ calls at a value of 0.85, and then required that > 30% of nucleotides along a molecule fell above this threshold value.

### Processed data analyses

All processed data analyses and associated scripts will be made available at https://github.com/RamaniLab/.

#### Global accessibility fold change analyses

For genome-wide data, we calculated accessibility fold change as the mean of HMM-derived accessibility for all BrdU^(+)^ molecules, divided by the mean of HMM-derived accessibility for an equal number of randomly sampled unlabeled molecules. These were calculated for each biological replicate and plotted.

#### Autocorrelation and clustering analyses

All autocorrelation analyses were performed as in ^20,28^; briefly, we computed the autocorrelation for the first 1 kb of all BrdU^(+)^ molecules >= 1 kb in length, as well as an equal number of randomly sampled steady state molecules. We then clustered these via leiden clustering, using the scanpy package. Leiden clustering was performed using resolution = 0.4; clusters were then filtered on the basis of size, such that all clusters that collectively summed up to < 10% of the total dataset were removed. Following cluster definition and filtering, we performed a series of Fisher’s exact tests to determine enrichment and depletion of clusters (*i.e.* ‘fiber types’) at each labeling time. To ascertain significance of enrichment while accounting for multiple hypothesis testing, we then converted all *p*-values into Storey *q*-values and drew an FDR cutoff of 5%.

#### Correlating sequence and signal

To correlate signal and sequence for irregular and regular BrdU^(+)^ and steady-state fibers, we extracted the first 1000 nucleotides of each read (same sequence used for autocorrelation calculations), and calculated the proportion of AA / TT/ AT / TA dinucleotides in a rolling 20 nucleotide window across the sequence. We then correlated that against the accessibility signal for that same region, for each molecule. To define IR versus non-IR (for K562 and mESC), we re-clustered BrdU^(+)^ molecules as above, and manually annotated clusters by visually inspecting average autocorrelations for each cluster. We note that the cluster definitions used for these analyses differ very slightly from those shown in Figure 3, due to differences in the random sampling of unlabeled fibers for each time point.

#### Epigenome-specific analyses

Custom scripts were used to isolate reads aligning to histone modification domains (derived from ENCODE), repeat sequences (see below), or rDNA (obtained by aligning CCS reads directly against a single copy of the murine rDNA repeat). Footprint sizes for these repeats from BrdU^(+)^ and unlabeled molecules were then plotted for each labeling time.

#### TF accessibility fold change analyses

Previously-written ^20,31^ custom scripts were used to isolate reads harboring ChIP-seq backed binding sites for CTCF, NRF1, REST, and SOX2, and TSSs, as well as control reads harboring random motif matches for each factor (lowly expressed TSSs for TSS) ^53^. Reads were filtered such that the motif was flanked by at least 750 bp on either side (1.5 kb window surrounding the motif). We then computed the mean accessibility of the 150 central bp of this window in BrdU^(+)^ versus an equal number of randomly sampled unlabeled molecules, for each timepoint. To determine whether TF accessibility fold changes between *bona fide* binding sites and random motif matches were significantly different, this process was repeated *n =* 1000 times per biological replicate, and a *p*-value was estimated via one-sided permutation test. All boxplots visualize the median and interquartile range of each distribution.

#### TF accessibility visualization

We plotted the mean signal for BrdU^(+)^ molecules (centered at each motif) in blue, against the mean signal for unlabeled molecules in red. For unlabeled molecules, we estimated a standard deviation of the mean signal (grey ribbon) by randomly sampling *n* = 100 equivalent numbers of unlabeled molecules for each labeling time.

#### TF accessibility clustering

We employed leiden clustering (resolution = 0.2) to cluster CTCF and transcribed TSSs. We then used Fisher’s exact tests to ascertain significant enrichment or depletion (*q*-value corrected, *q* < 0.05) of each cluster as a function of labeling time.

#### Satellite sequence analyses

Mouse minor satellite (centromeric), major satellite (pericentromeric), and telomeric reads were identified directly from PacBio CCS reads as previously described ^20^.

## DATA AVAILABILITY

Raw and processed data will be made available at GEO accession GSEXXXX.

## Supporting information

Supplemental tables referenced in Ostrowski, Yang, et al. manuscript

## ACKNOWLEDGEMENTS

The authors thank Daniele Canzio (UCSF), Hiten Madhani (UCSF), and Geeta Narlikar (UCSF) for helpful discussions and comments on the manuscript.

